# Group-common and individual-specific effects of structure-function coupling in human brain networks with graph neural networks

**DOI:** 10.1101/2023.11.22.568257

**Authors:** Peiyu Chen, Hang Yang, Xin Zheng, Hai Jia, Jiachang Hao, Xiaoyu Xu, Chao Li, Xiaosong He, Runsen Chen, Tatsuo S. Okubo, Zaixu Cui

**Affiliations:** Beijing Institute for Brain Research, Chinese Academy of Medical Sciences & Peking Union Medical College, Beijing, 102206, China; Chinese Institute for Brain Research, Beijing; Beijing, 102206, China; Department of Biomedical Engineering, Vanderbilt University, Nashville, TN 37235, USA; State Key Laboratory of Cognitive Neuroscience and Learning, Beijing Normal University, Beijing, 100091, China; Department of Applied Mathematics and Theoretical Physics, Department of Clinical Neurosciences, University of Cambridge, Cambridge, CB3 0WA, UK; Department of Psychology, School of Humanities and Social Sciences, University of Science and Technology of China, Hefei, Anhui, 230026, China; Vanke School of Public Health, Tsinghua University, Beijing, China

**Keywords:** diffusion MRI, functional MRI, structure-function coupling, graph neural networks, individual effects, sensorimotor-association cortical axis

## Abstract

The human cerebral cortex is organized into functionally segregated but synchronized regions bridged by the structural connectivity of white matter pathways. While structure-function coupling has been implicated in cognitive development and neuropsychiatric disorders, it remains unclear to what extent the structure-function coupling reflects a group-common characteristic or varies across individuals, at both the global and regional brain levels. By leveraging two independent, high-quality datasets, we found that the graph neural network accurately predicted unseen individuals’ functional connectivity from structural connectivity, reflecting a strong structure-function coupling. This coupling was primarily driven by network topology and was substantially stronger than that of the correlation approaches. Moreover, we observed that structure-function coupling was dominated by group-common effects, with subtle yet significant individual-specific effects. The regional group and individual effects of coupling were hierarchically organized across the cortex along a sensorimotor-association axis, with lower group and higher individual effects in association cortices. These findings emphasize the importance of considering both group and individual effects in understanding cortical structure-function coupling, suggesting insights into interpreting individual differences of the coupling and informing connectivity-guided therapeutics.

## 1. INTRODUCTION

The human cerebral cortex is organized into functionally segregated neuronal populations connected by anatomical pathways. White matter fiber tracts form a connectome of structural connectivity at the macroscale (Sporns, 2011). This structural connectome exhibits a complex network topology characterized by non-random properties, including small-world architecture (Bassett & Bullmore, 2006), segregated communities (Sporns & Betzel, 2016), and a core of densely interconnected hubs (van den Heuvel & Sporns, 2013). These topological patterns support the communication dynamics of structural networks and coordinate the temporal synchronization of neural activity, termed functional connectivity, between cortical regions (Avena-Koenigsberger et al., 2017; Lynn & Bassett, 2019; Seguin, Sporns, et al., 2023; Suárez et al., 2020). Understanding how structural connectivity shapes functional connectivity patterns is central to neuroscientific research.

Convergent evidence from multiple independent studies suggests a reliable coupling between structural and functional connectivity at both global and regional brain levels, using non-invasive magnetic resonance imaging (MRI) techniques and invasive recordings (Baum et al., 2020; Betzel et al., 2019; Deco et al., 2013; Demirtas et al., 2019; Deslauriers-Gauthier et al., 2022; Gu et al., 2021; Honey et al., 2009; Medaglia et al., 2018; Misic et al., 2016; Misic et al., 2015; Sarwar et al., 2021; Seguin, Jedynak, et al., 2023; Valk et al., 2022; Vazquez-Rodriguez et al., 2019; Zamani Esfahlani et al., 2022). The structure-function coupling is heterogeneously distributed across the cerebral cortex, exhibiting higher coupling in the primary sensorimotor cortex and lower coupling in the higher-order association cortex (Baum et al., 2020; Gu et al., 2021; Vazquez-Rodriguez et al., 2019). This spatial distribution pattern aligns with the sensorimotor-association cortical hierarchy of cytoarchitectonic structure, functional specialization, and evolutionary expansion (Baum et al., 2020; Sydnor et al., 2021; Vazquez-Rodriguez et al., 2019). The structure-function coupling shows a developmental increase at regions of association cortex during the adolescence with the most prominent effects localized in the default mode network (Baum et al., 2020). In contrast, the highly evolutionarily conserved sensorimotor regions exhibit age-related decreases in structure-function coupling throughout adolescence (Baum et al., 2020) and the whole lifespan (Zamani Esfahlani et al., 2022). Moreover, higher structure-function coupling has been related to better performance in executive function (Baum et al., 2020; Medaglia et al., 2018), and the abnormal pattern of the coupling is associated with a wide range of psychiatric and neurological disorders, such as major depressive disorder (Jiang et al., 2019), bipolar disorder (Jiang et al., 2020), attention deficit hyperactivity disorder (Soman et al., 2023), and Parkinson’s disease (Zarkali et al., 2021).

While structure-function coupling has been extensively implicated in development, cognition, and clinical outcomes, few studies have explicitly examined the extent to which structure-function coupling reflects a group-common characteristic or varies across individuals. Prior studies have indicated that functional connectivity is dominated by both stable group and individual factors (Gratton et al., 2018), while structural connectivity exhibits less variability across participants (Zimmermann et al., 2019). This leads to a hypothesis that structure-function coupling may primarily reflect group characteristics rather than individual differences. One prior study examined the presence of individual-specific effects in structure-function coupling through Pearson correlation and reported conflicting findings (Zimmermann et al., 2019). These discrepancies could arise from the application of the correlation approach, which solely accounts for the coupling of direct structural connectivity while disregarding indirect functional communications. To date, the magnitudes of group-common and individual-specific effects of structure-function coupling remains unclear. Additionally, no study has evaluated group and individual coupling at the regional level or explored the cortical organization of these regional group and individual structure-function couplings.

To fill these gaps, we used graph neural networks (GNNs) to investigate the magnitude of group-common and individual-specific effects of the structure-function coupling at both global and regional levels. GNNs can capture indirect and non-linear inter-regional functional communication on the structural connectome (Neudorf et al., 2022; Zhou et al., 2020), whereas traditional correlation (Baum et al., 2020; Honey et al., 2009) or multilinear regression (Vazquez-Rodriguez et al., 2019) approaches only account for direct structural connections when estimating structure-function coupling. We tested the following three interrelated hypotheses: First, we hypothesized that we would replicate prior findings (Neudorf et al., 2022) that the GNN accurately predicts functional connectivity from structural networks. We also hypothesized that this prediction would primarily be determined by higher-order network topology, as GNNs inherently capture the graph topology between network nodes (Neudorf et al., 2022; Zhou et al., 2020). Second, we hypothesized that structure-function coupling would mainly reflect group-common characteristics (Zimmermann et al., 2019); however, there would still exist a significant amount of individual effect, given the implications of structure-function coupling in human individual differences (Baum et al., 2020; Zamani Esfahlani et al., 2022). Furthermore, we predicted that GNNs might better identify individual coupling effects compared to traditional correlation methods, since GNNs take into account indirect functional communications. Third, we hypothesized that both group and individual effects at the regional level would be distributed across the cortex along the canonical sensorimotor-association cortical axis (Sydnor et al., 2021), which is a convergent organizing principle unifying diverse neurobiological properties. Specifically, given that sensorimotor cortices exhibit less inter-individual variability in both structural and functional connectivity (Gratton et al., 2018; Huang et al., 2023; Mueller et al., 2013), we hypothesized that the individual-specific effects of structure-function coupling would also be lower in the sensorimotor cortices. In contrast, the association cortices demonstrate greater inter-individual variability in both structural and functional connectivity (Gratton et al., 2018; Huang et al., 2023; Mueller et al., 2013). Therefore, we expected that the individual-specific effects would be stronger and group-common effects would be lower in the higher-order association cortex.

## 2. METHODS

### 2.1. Study cohort

#### 2.1.1. HCP young adult (HCP-YA) dataset

We acquired multi-modal neuroimaging data from 339 unrelated participants (156 males, aged 22–37) from the HCP young adult (HCP-YA) dataset (release S900), including T1-weighted structural, resting-state functional MRI (fMRI) and diffusion MRI (Van Essen et al., 2013). All imaging data were acquired by a multiband sequence on a Siemens 3T Skyra scanner. Two resting-state fMRI sessions, with two runs in each session (left-right and right-left phase-encoding), were acquired for each participant with a resolution of 2 mm isotropic. Each resting-state run comprised 1,200 frames for approximately 15 min in length. For diffusion MRI, data were acquired in two runs with opposite phase-encoding directions for each participant. Each run included 270 non-collinear directions with 3 non-zero shells (b = 1000, 2000, and 3000 s/mm^2^). Other details regarding the HCP-YA dataset and MRI acquisition parameters have been described in prior study (Van Essen et al., 2013).

#### 2.1.2. HCP development (HCP-D) dataset

This study also comprised 633 participants (294 males, aged 8–21) obtained from the HCP-development (HCP-D) dataset (Release 2.0) (Somerville et al., 2018). All data were collected using a multiband EPI sequence on a 3T Siemens Prisma scanner. Two resting-state fMRI sessions were acquired for each participant, with two runs in each session, using anterior-posterior (AP) and posterior-anterior (PA) phase-encoding, respectively. Each resting-state run was approximately 6.5 min with 488 frames. Diffusion MRI data included two sessions, each with two non-zero shells (b = 1500, 3000 s/mm^2^) and 185 diffusion-weighted directions. Further details about the HCP-D dataset have been described in previous study (Somerville et al., 2018).

### 2.2. MRI data processing

#### 2.2.1. Structural and functional MRI data processing

Minimally preprocessed T1-weighted structural and functional MRI data were acquired from the HCP-D and HCP-YA datasets (Glasser et al., 2013). The HCP minimal preprocessing pipelines used the tools FreeSurfer (Fischl, 2012), FSL (Smith et al., 2004), and Connectome Workbench (Marcus et al., 2011). Briefly, structural MRI data were corrected for intensity non-uniformity, skull-stripped, and then used for cortical surface reconstruction. Volume-based structural images were segmented into cerebrospinal fluid (CSF), white matter, and gray matter, and spatially normalized to the standard MNI space. Functional MRI data were preprocessed with slice-timing, motion and distortion correction, co-registration to structural data, normalization to MNI space, and projection to cortical surface. Functional timeseries were resampled to FreeSurfer’s fsaverage space, and grayordinates files containing 91k samples were generated (Glasser et al., 2013).

We then followed the post-processed protocols of eXtensible Connectivity Pipelines (XCP-D; https://xcp-d.readthedocs.io/en/latest/) (Mehta et al., 2024). We regressed out 36 nuisance regressors from the BOLD time series, including six motion parameters, global signal, mean white matter signal, and mean CSF signal, along with their temporal derivatives, quadratic terms, and quadratic derivatives (Ciric et al., 2017). Residual timeseries were then band-pass filtered (0.01–0.08 Hz). We quantified head motion by calculating Jenkinson’s framewise displacement (FD) (Jenkinson et al., 2002), and we further reduced the potential effects of head motion by excluding subjects based on two criteria (Faskowitz et al., 2020). First, we dropped those fMRI runs that included more than 25% of the frames with FD > 0.2 mm. Second, we calculated the mean FD distribution for each fMRI run by pooling frames of all participants. Then, we derived the third quartile (Q3) and interquartile range (IQR) of this distribution. Runs with mean FD greater than “Q3+1.5×IQR” were excluded. We excluded 111 HCP-D and 42 HCP-YA participants based on these two criteria. Additionally, seven participants from the HCP-D were excluded because of less than four resting-state fMRI runs. Consequently, 514 participants from the HCP-D and 297 participants from the HCP-YA datasets were included in subsequent analyses.

#### 2.2.2. Functional connectivity construction with functional MRI

Functional connectivity (FC) refers to the temporal correlation of functional MRI signals. For HCP-YA and HCP-D datasets, we constructed FC by the following procedures. First, we extracted regional BOLD timeseries based on the prior Schaefer parcellation with 400 parcels (Schaefer et al., 2018). In the atlas, these 400 parcels have been assigned to seven large-scale functional networks (Yeo et al., 2011), including the visual network (VIS), somatomotor network (SMN), dorsal attention network (DAN), ventral attention network (VAN), frontoparietal network (FPN), limbic network (LIM), and default mode network (DMN). Next, FC was calculated as the Pearson correlation coefficient between each pair of regional BOLD timeseries, resulting in a 400×400 symmetrical FC matrix for each participant. We then applied Fisher’s z-transformation to each FC value in the matrix.

#### 2.2.3. Diffusion MRI data processing

We used minimally preprocessed diffusion MRI data from the HCP-YA dataset (Van Essen et al., 2013). The minimal preprocessing pipeline comprises b0 image intensity normalization across runs, EPI distortion correction, eddy current and motion correction, gradient nonlinearity correction, and registration to the native structural space (1.25 mm). The processed diffusion MRI data were further corrected for B1 field inhomogeneity using MRtrix3 (https://www.mrtrix.org/) (Tournier et al., 2019). Diffusion MRI data from HCP-D were preprocessed by QSIPrep (https://qsiprep.readthedocs.io/), an integrative platform for preprocessing and reconstructing diffusion MRI data (Cieslak et al., 2021), including tools from MRtrix3. Prior to preprocessing, we concatenated the two AP runs and the two PA runs, respectively, and selected the frames with b-value < 100 s/mm^2^ as the b0 image. Next, we applied MP-PCA denoising, Gibbs unringing, and B1 field inhomogeneity correction through MRtrix3’s *dwidenoise* (Veraart et al., 2016), *mrdegibbs* (Kellner et al., 2016), and *dwibiascorrect* (Tustison et al., 2010) functions. FSL’s eddy was then used for head motion correction and Eddy current correction (Andersson et al., 2016). Finally, the preprocessed DWI timeseries was resampled to ACPC space at a resolution of 1.5 mm isotropic. Particularly, a total of 15 participants from the HCP-D were excluded from subsequent structural connectivity analyses due to incomplete DWI data (8 participants), tissue segmentation failure (5 participants), or the identification of isolated regions after tractography (2 participants). Therefore, a total of 297 participants from HCP-YA (137 males, aged 22–36) and 499 participants from HCP-D (237 males, aged 8–21) were analyzed in subsequent structure-function coupling analyses.

#### 2.2.4. White matter structural network construction with diffusion MRI

We reconstructed whole-brain white matter tracts from preprocessed diffusion MRI data to construct the structural connectivity (SC). Reconstruction was conducted by the *mrtrix_multishell_msmt_ACT-hsvs* method in MRtrix3 (Tournier et al., 2019), which implements a multi-shell and multi-tissue constrained spherical deconvolution (CSD) to estimate the fiber orientation distribution (FOD) of each voxel (Jeurissen et al., 2014). Then, we followed the anatomically constrained tractography (ACT) framework to improve the biological accuracy of fiber reconstruction (Smith et al., 2012). This tractography was performed by *tckgen*, which generates 40 million streamlines (length range from 30 to 250 mm, FOD power = 0.33) via a refined probabilistic streamlines tractography (iFOD2) based on the second-order integration over FOD. The streamlines were filtered from the tractogram based on the spherical deconvolution of the diffusion signal. We estimated the streamline weights using the command *tcksift2* (Smith et al., 2015). Next, the SC matrix was constructed by *tck2connectome* based on the Schaefer-400 atlas for each participant. The edge weight of SC indicates the number of streamlines connecting two regions. For each connection, we normalized the edge weight by dividing the average volume of the corresponding two regions (Hagmann et al., 2008). Furthermore, the edge weight of the SC matrix was log-transformed, which is commonly used to shift the non-Gaussian SC distribution to the Gaussian distribution in previous studies (Suárez et al., 2020). Finally, we obtained a 400×400 symmetric SC matrix for each participant. To address potential issues with spurious SC edges arising from probabilistic tractography (Roberts et al., 2017), we adopted a group-consistency thresholding approach as in previous studies (Baum et al., 2020; Ge et al., 2023). We calculated the coefficient of variation (CV) for edge weight across participants and subsequently applied a threshold to individual structural connectivity matrices at the 75th percentile for edge weight CV. The top quartile of inconsistent connections was removed.

### 2.3. Predicting functional connectivity from structural connectivity using graph neural networks

#### 2.3.1. Graph convolutional network

The main component of our graph neural network is the graph convolutional network (GCN) (Kipf & Welling, 2017). GCN generates node embeddings, representing the neighboring topological structure for each node. We denoted the node embedding generated from *l*^th^ layer GCN as 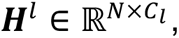 where *l* ∈ {1, …, *L*} represents the layer index and *C*_*l*_ represents the embedding dimension of layer *l*. *H*^0^ is the exception representing the dimension of the input features of GCN. The input features are one-hot vectors that indicate the brain region of each node. Through message passing, the node embedding from the *l*^th^ layer will pass along the network connectivity, denoted its adjacency matrix as ***A***, towards their neighboring nodes, convoluted by a GCN layer to get the node embedding of the (*l* + 1)^th^ layer. In our work, ***A*** is the adjacency matrix of SC, i.e., ***A*** ∈ ℝ^*K*×*K*^. *K* = 400 is the number of nodes for the Schaefer-400 atlas. Therefore, the update equation of the node embedding from the *l*^th^ layer to the (*l* + 1)^th^ layer is denoted as:

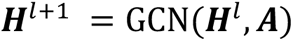

With *L* layers in total, this message passing allows each node to embed its L-hop topological neighborhood. The concrete equation is formulated as:

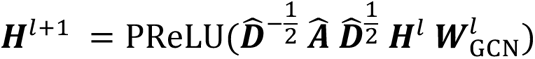

where 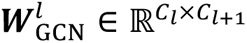 is the trainable weight at the *l*^th^ layer. 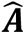 is the adjacency matrix with self-loop added, i.e., 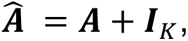 where 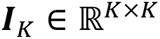 is the K-dimension identity matrix. 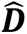 is the diagonal matrix, with the diagonal value 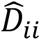 equals to the nodal strength of node *i* in ***A***. As such, 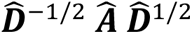 is the symmetric normalized Laplacian of the SC. We adopted the PReLU (Parametric Rectified Linear Unit) as the activation function due to its superior performance (You et al., 2021):

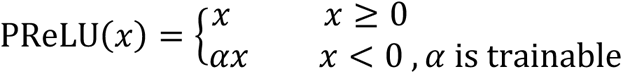

#### 2.3.2. Multilayer perceptron

Subsequently, we used a pair of node embeddings as input to predict the functional connectivity between them by a two-layer multi-layer perceptron (MLP) with ReLU (Rectified Linear Unit) as the activation function (i.e., ReLU(*x*) = max(0, *x*)) after the first output layer. No activation function was used for the second layer. Specifically, for two node embeddings of node *i* and *j*, represented by ℎ*_i_*, ℎ*_j_* ∈ *H*, the predicted FC, termed pFC*_ij_* ∈ **pFC**, between them is calculated as:

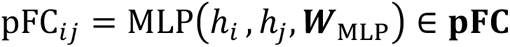

#### 2.3.3. Model training and validation

To train our graph neural network model, we minimized the mean square error between the predicted FC and empirical FC with L2-regularizaion:

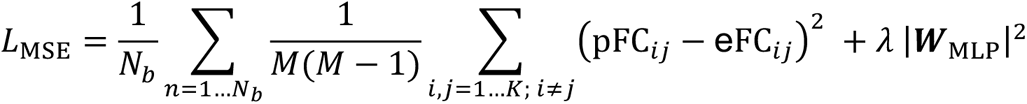

where eFC*_ij_* ∈ **eFC** is the empirical FC between node *i* and *j*. We adopted L2-regularization on the MLP’s weights with regularization parameter *λ*. We trained the model through batched gradient descent with a batch of *N*_*b*_subjects and the Adam optimizer (Kingma & Ba, 2017). In addition, since FC is a symmetric matrix, we employed the (pFC*_ij_* + pFC*_ji_*)/2 as the final predicted FC between node *i* and *j* for testing and further analysis. For both datasets, we randomly split 50% of the subjects as the training set, with the remaining 50% for testing and subsequent structure-function coupling analysis. By grid search based on the model’s performance for 5-fold cross validation on the training set, we set the hyperparameter as: layer number (L = 2), dimension for GCN (400×256×256), dimension for MLP (512×64×1), batch size (2), learning rate (0.001), regularization parameter (*λ* = 0.0001), and training epochs (400). The graph network model was built and trained using Pytorch and PyTorch Geometric. More details about implementations could be found in the code repository at https://github.com/CuiLabCIBR/GNN_SC_FC.

### 2.4. Structure-function coupling

#### 2.4.1. Structure-function coupling measured using correlation and GNNs

In this work, we evaluated the structure-function coupling using both a correlation approach and our GNN framework. The correlation approach assesses the Pearson correlation between SC and FC, focusing solely on direct connections with nonzero edge strength. However, functional connectivity can emerge from polysynaptic communication on structural connectome (Avena-Koenigsberger et al., 2017; Seguin, Sporns, et al., 2023). The GNN has the capability to capture both direct and indirect functional communications based on the sparse SC. For each dataset, all participants were split into two subsets with one used as training and another used as test set. The GNN model trained on all participants in the training set was used to predict the FC for the testing participants with their SC as input. The structure-function coupling was calculated as the Pearson correlation between the predicted and empirical FC for each participant in the test set. At the group level, we used the model from training data and all testing participants’ average SC as input to predict the average FC. We calculated the Pearson correlation between the predicted and empirical average FC as the structure-function coupling at the group level.

We also estimated the structure-function coupling at both the whole-brain and regional levels. At the whole-brain level, we extracted the upper triangle elements (79,800 unique edges) from either the 400×400 SC matrix or the predicted FC matrix, creating a structural profile vector. Simultaneously, we extracted the upper triangle elements from the empirical FC matrix, forming a functional profile vector. We calculated the Pearson correlation between the structural profile and functional profile across all connections at the global level. We reported across-participant mean and standard deviation (SD) of individual structure-function coupling as mean ± SD. At the regional level, we first defined the regional structural and functional profiles. For the *i*^th^ cortical region, we defined its structural or functional profile as the connections to all other cortical regions, which were represented by the *i*^th^ row (399×1) from the connection matrix. The regional structure-function coupling was determined by calculating the Pearson correlation between the structural profile and functional profile of each cortical region, enabling us to generate a cortical map of regional structure-function coupling for both the correlation approach and the GNN model. Notably, all participants were involved and analyzed in the correlation-based coupling, whereas the structure-function coupling based on the GNN model was derived from the test set, comprising 50% of the participants.

#### 2.4.2. Calculating group-common and individual-specific effects in structure-function coupling

A previous study indicated that the individual FC is contributed by both group-common and stable individual-specific factors (Gratton et al., 2018). Following a similar approach, we aimed to distinguish between group-common and individual-specific effects in structure-function coupling using both the correlation approach and the GNN model. To achieve this, we computed the structure-function coupling both within-individual and between each pair of different participants, resulting in a participant-by-participant asymmetrical matrix of structure-function coupling (**Fig. 3a**). In this matrix, the element in the *i*^th^ row (SC or predicted FC from subject *i*) and *j*^th^ column (FC from subject *j*), termed as cp*_i,j_*, represents the structure-function coupling between the *i*^th^ and *j*^th^ participants. The diagonal elements of the matrix measure the structure-function coupling within individual, while the off-diagonal elements estimate the coupling between one individual’s SC and another individual’s FC. We referred to the within-individual coupling and between-individual coupling as “matched coupling” and “mismatched coupling”, respectively. In this study, we treated the averaged between-individual coupling as a group-common variance in the coupling across populations. This strategy has been used in a prior study of effects in function networks (Gratton et al., 2018). We supposed that the total effects of within-individual coupling encompass both group-common and individual-specific effects. Therefore, we defined the group-common and individual-specific effects of structure-function coupling as follows:

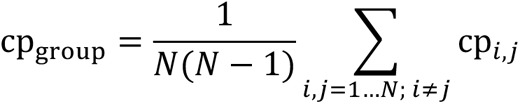

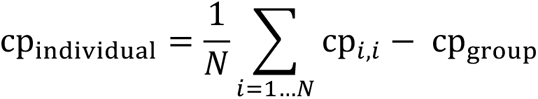

where *N* corresponds to the number of participants. Given our finding that individual structure-function coupling is primarily dominated by the group-common effect, with minor individual-specific effects (see **RESULTS**), we proceeded to assess the statistical significance of these individual effects. For this purpose, we defined the matched coupling and mismatched coupling for each subject (**Fig. 3d**). Specifically, the mismatched coupling of subject *i* was defined by the averaged coupling from all elements in the *i*^th^ row (SC or predicted FC from subject *i*, FC from other subjects) and *i*^th^ column (FC from subject *i*, SC or predicted FC from other subjects), except the within-individual coupling. The matched and mismatched coupling for each subject *i* is defined as:

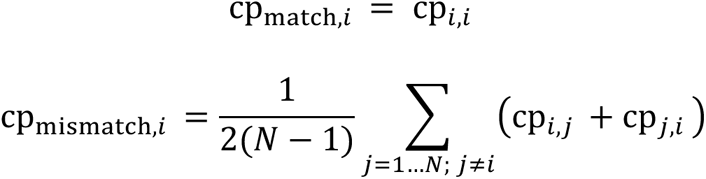

We considered the individual-specific coupling effects significant if matched coupling were statistically higher than mismatched coupling across participants. We conducted a two-sided paired t-test to examine the differences between matched coupling and mismatched coupling across all participants.

These analyses of group and individual effects in the coupling were first performed at the global level and then applied to each cortical region. It is noteworthy that only nonzero edges were examined in the correlation approach for both whole-brain and regional analyses.

#### 2.4.3. Alignment between cortical maps of group and individual effects of structure-function coupling and the sensorimotor-association cortical axis

A recent study proposed a canonical sensorimotor-association cortical axis to characterize a dominant pattern across the human cortex, along which diverse neurobiological properties progressively change in a continuous order (Sydnor et al., 2021). Previous studies have shown that the cortical distribution of structure-function coupling share a similar pattern with sensorimotor-association cortical axis across the cortex (Baum et al., 2020; Vazquez-Rodriguez et al., 2019). Here, we replicated this finding and further investigated how group-common and individual-specific coupling aligns with the sensorimotor-association cortical axis, respectively. We acquired a cortical map of the sensorimotor-association axis parcellated with the Schaefer-400 atlas from Sydnor et al. (Sydnor et al., 2021). Cortical regions were continuously ranked along this axis, with the primary sensorimotor cortices representing the lowest ranks and the higher-order association cortices representing the highest ranks. We first evaluated the Spearman’s rank correlation between structure-function coupling and sensorimotor-association cortical ranks across all regions for both correlation- and GNN-based coupling. We further calculated Spearman’s rank correlation to assess the alignment between the sensorimotor-association cortical axis and regional group-common effects as well as individual-specific effects across all cortical regions for both coupling approaches.

### 2.5. Null models

#### 2.5.1. Rewiring networks

To evaluate the contribution of network topology in GNN-based structure-function coupling, we utilized the Maslov-Sneppen rewiring algorithm to rewire individual SC (Maslov & Sneppen, 2002). This process retained nodal degree and strength distribution while randomizing network topological structure. This algorithm was implemented via the Python version of the Brain Connectivity Toolbox (https://github.com/aestrivex/bctpy) (Rubinov & Sporns, 2010). We aimed to evaluate how much the GNN model could learn from the higher-order topology, so we only rewired SC in training data to acquire a topological-null GNN model. We did not rewire the SC in the test set. We next used this topological-null GNN model to predict FC from SC and re-evaluated the Pearson correlation between the predicted FC and FC for each participant in the test set.

#### 2.5.2. Spin test

We employed the spin test (Alexander-Bloch et al., 2018) to evaluate the significance of the spatial correspondence between group-common and individual-specific effects of structure-function coupling and sensorimotor-association cortical axis. Particularly, the spin test generated a null distribution by randomly rotating brain maps while maintaining the original spatial covariance structure (Alexander-Bloch et al., 2018). This approach projected the group and individual effects onto a spherical cortical surface of the FreeSurfer’s fsaverage space and then randomly rotated 10,000 times to generate a list of rotated maps. Next, we calculated Spearman’s rank correlation between each rotated map and the sensorimotor-association axis map to construct a null distribution. The *P* value (*P*_spin_) was determined by calculating the ratio of instances whose null correlations exceeded the empirical correlation coefficients.

### 2.6. Sensitivity analyses

To validate our results and ensure their robustness to train-test splits, we implemented repeated 2-fold cross-validations for our GNN models, while keeping the architecture and the hyperparameters the same as in the main analyses. Specifically, for each dataset, participants were randomly divided into two equally sized folds, with one using as the training set and the other served as the test set. This process was repeated after reversing the role of training set and test set, ensuring that each fold was used for both training and testing. During each of the two iterations, the GNN model was trained on the training data and evaluated on the test data, followed by structure-function coupling analyses on the test set. This 2-fold cross-validation was repeated 100 times to ensure robustness to train-test split.

As individual-specific effects are derived by subtracting the group-common effects from the total effects of the coupling, larger total effects might numerically lead to larger individual-specific effects. To address this potential scaling issue, we defined the “normalized individual effect” by dividing the individual effect by the total regional structure-function coupling, representing the proportion of the individual effect to the total effect. Subsequently, we re-evaluated the alignment between the normalized individual effects and the ranks of the sensorimotor-association cortical axis for the GNN model.

Finally, we assessed the robustness of our findings to the variation of cortical parcellations with Schaefer atlas of 200 cortical regions (Schaefer et al., 2018). We constructed the connectome of FC and SC for each participant using the Shcaefer-200 atlas and retrained the GNN models. We repeated all our primary analyses using these networks, including quantifying both the global and regional group-common and individual-specific effects of structure-function coupling and evaluating the alignment between regional couplings and sensorimotor-association cortical axis.

## 3. RESULTS

We utilized two independent datasets, namely the Human Connectome Project (HCP)-young adult (HCP-YA, *n* = 297, 137 males, aged 22–36 years) (Van Essen et al., 2013) and HCP-development (HCP-D, *n* = 499, 237 males, aged 8–21 years) (Somerville et al., 2018), to study structure-function coupling. As illustrated in **Fig. 1a**, we first constructed the connectome of functional connectivity (FC) and structural connectivity (SC) for each participant using a Schaefer cortical parcellation atlas of 400 regions (Schaefer et al., 2018). Based on the resting-state functional MRI (fMRI) data, FC was defined as the Pearson correlation coefficient between each pair of regional time series, resulting in a 400×400 symmetrical FC matrix for each participant. The matrix consisted of 79,800 unique elements, with each element denoted as an “edge” connecting two cortical regions. Fisher *r*-to-*z* was used to improve the normality of the FC edge strength. Using diffusion MRI data, we reconstructed whole-brain white matter tracts of individual participants using probabilistic fiber tractography with multi-shell, multi-tissue constrained spherical deconvolution (CSD) (Jeurissen et al., 2014). Anatomically constrained tractography (ACT) (Smith et al., 2012) and spherical deconvolution informed filtering of tractograms (SIFT) (Smith et al., 2015) have been applied to improve the biological accuracy of fiber reconstruction. For each participant, we quantified the number of streamlines connecting each pair of cortical regions using the Schaefer atlas to construct a structural connectome of the streamline counts. The edge weights of the SC matrices were log transformed.

**Fig. 1.**
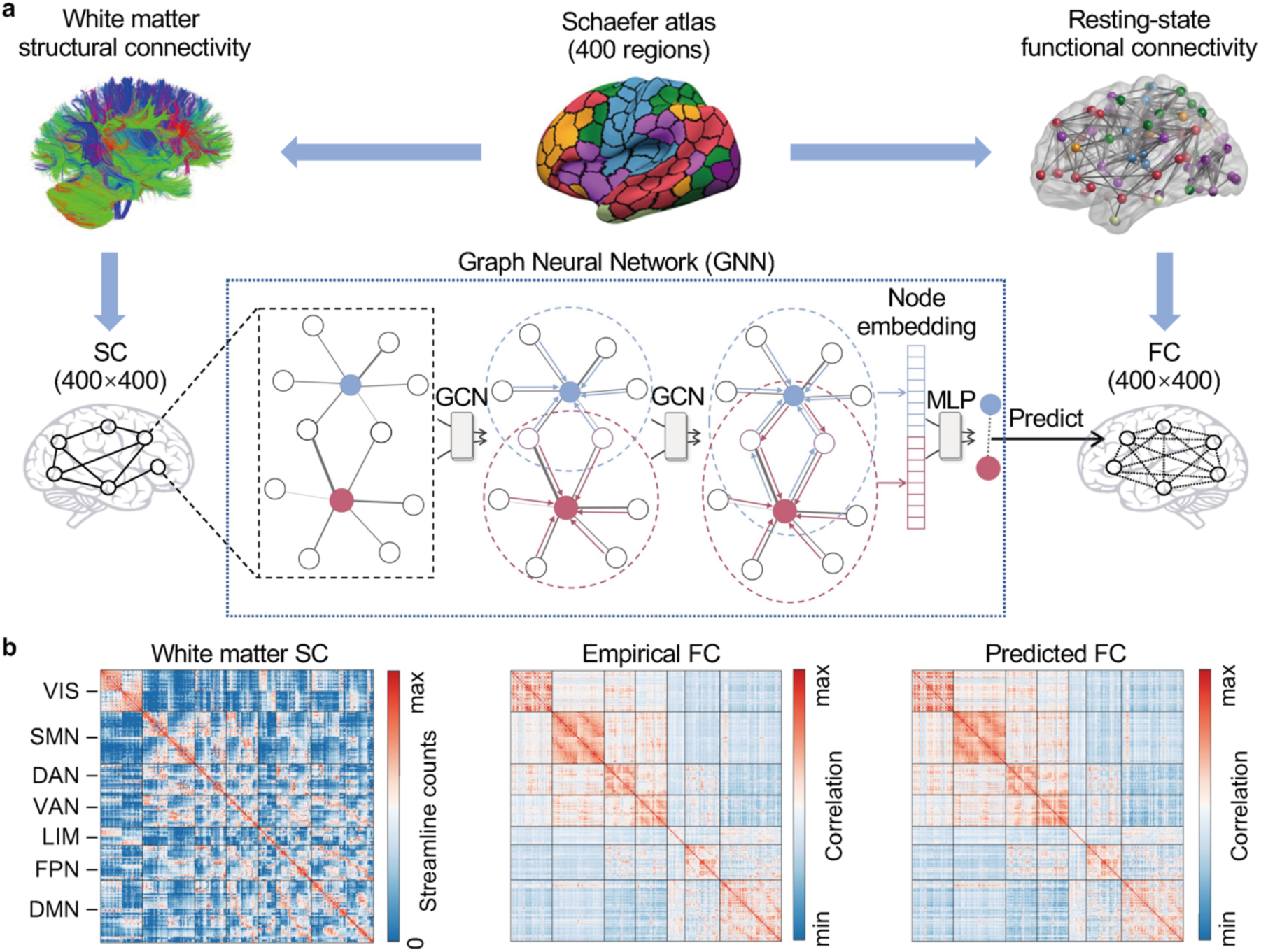
The graph neural network (GNN) predicts functional connectivity from structural connectivity. **a**, For each individual, the white matter structural connectivity (SC) and resting-state functional connectivity (FC) were constructed using the Schaefer-400 atlas. Both SC and FC can be characterized as a 400×400 matrix. We utilized a GNN model to predict FC from SC. Specifically, the GNN employs a two-layer graph convolutional network (GCN) to get the node embedding for each node and uses a two-layer multilayer perceptron (MLP) to predict each FC edge from the corresponding two node embeddings. **b**, The group-averaged connectivity matrix of SC, empirical FC, and predicted FC in the HCP-YA dataset. Visual inspection suggests a highly consistent connectivity pattern between the empirical and predicted FC matrices. HCP-YA, Human Connectome Project (HCP)-young adult; VIS, visual network; SMN, somatomotor network; DAN, dorsal attention network; VAN, ventral attention network; LIM, limbic network; FPN, frontoparietal network; DMN, default mode network.

To fully investigate the indirect functional communication and nonlinear association between structural and functional connectomes, we used a graph neural network (GNN) framework to capture structure-function coupling by predicting the empirical FC from the SC (Hong et al., 2023; Neudorf et al., 2022; Sarwar et al., 2021). The GNN framework treats each FC edge as a higher-order communication through the SC between two corresponding nodes. Specifically, our GNN model employed a two-layer graph convolutional network (Kipf & Welling, 2017), which takes the SC as the input and aggregates the two-hop neighboring SC for each node to construct its node embeddings. Subsequently, a two-layer multilayer perceptron takes each pair of node embeddings as inputs to predict the connected FC (**Fig. 1a**). For both HCP-YA and HCP-D datasets, we randomly split the participants into two subsets, with one as the training set and the other as the test set. We trained a GNN model with the training data set to predict FC for each participant in the test set. We calculated the structure-function coupling for each participant in the test set with the Pearson correlation between the predicted and empirical FC. The subsequent GNN-based coupling analyses were restricted to the test data.

We presented the group-averaged connectivity matrices of SC, empirical FC, and predicted FC using the GNN in the HCP-YA dataset as examples (**Fig. 1b**). The connectivity matrix was organized based on the large-scale functional network affiliations of the 400 cortical regions to the Yeo 7 networks (Yeo et al., 2011), which consists of visual, somatomotor, dorsal attention, ventral attention, frontoparietal, limbic, and default mode networks. The predicted FC matrix was highly similar to the empirical FC matrix (**Fig. 1b**), suggesting the effectiveness of our GNN model. **Fig. S1** illustrates the empirical and predicted connectivity matrices of the HCP-D dataset.

### 3.1. GNN accurately captures the coupling between structural and functional network topology

Previous studies have consistently demonstrated robust coupling between SC and FC using correlation (Baum et al., 2020; Honey et al., 2009) or multilinear regression (Vazquez-Rodriguez et al., 2019) approaches. In this study, we replicated this finding in our datasets. We acquired group-averaged empirical FC and SC matrices and flattened the upper triangle elements into vectors for both matrices. Using Pearson correlation, we observed positive associations between FC and SC across edges with non-zero SC strength in both the HCP-YA (*r* = 0.396, *P* < 1×10^-16^; **Fig. 2a**) and HCP-D (*r* = 0.420, *P* < 1×10^-16^; **Fig. 2b**) datasets at the group level. At the individual level, we found that the correlation between FC and SC ranges from *r* = 0.215 to *r* = 0.389 (*r* = 0.297 ± 0.030 [mean ± SD]; **Fig. 2e**) across HCP-YA participants and ranges from *r* = 0.215 to *r* = 0.409 (*r* = 0.306 ± 0.028; **Fig. 2f**) across HCP-D participants. These results are consistent with the previously reported magnitudes of group-level structure-function coupling using the correlation approaches (Baum et al., 2020; Honey et al., 2009).

**Fig. 2.**
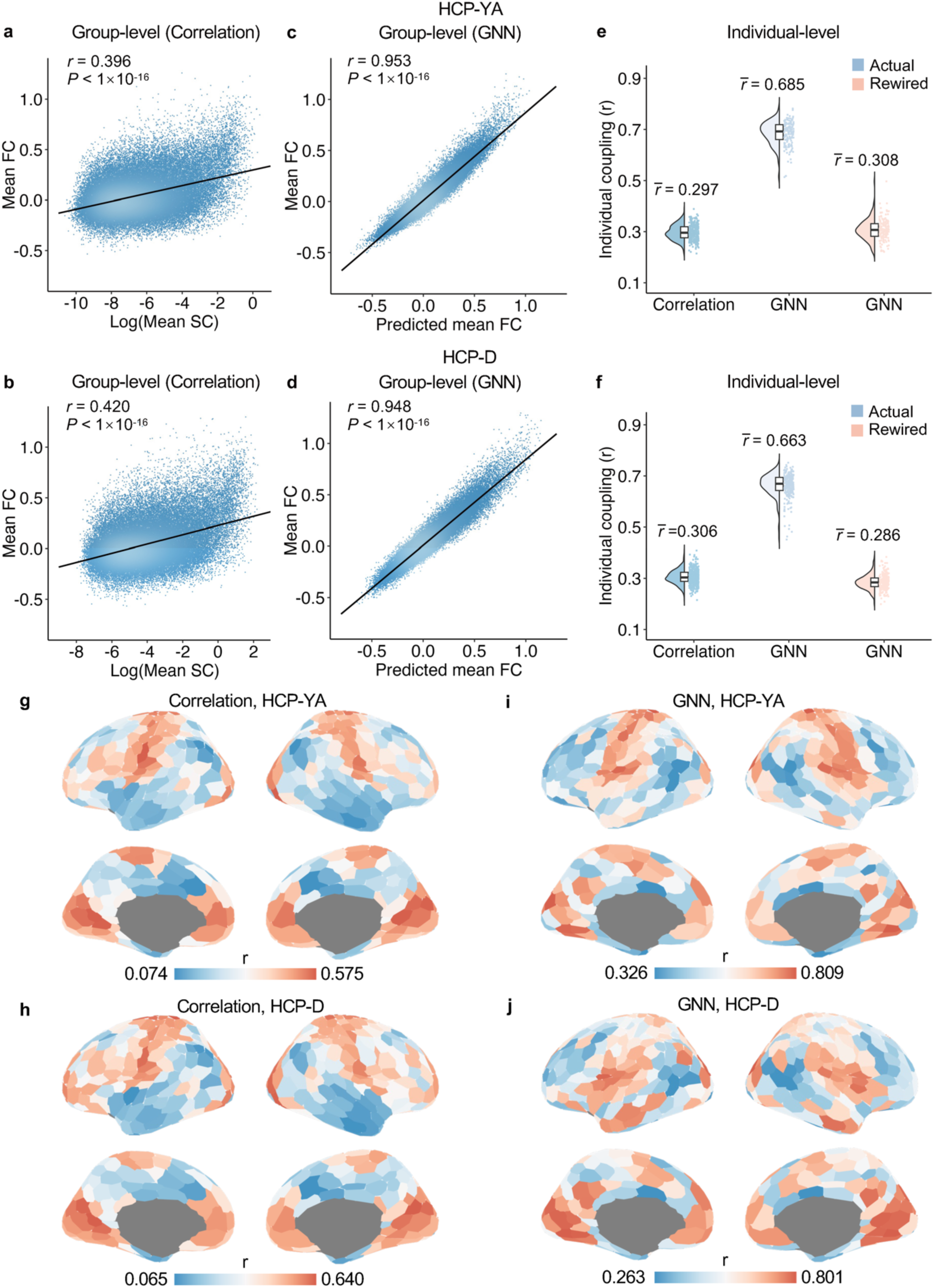
Structure-function coupling based on the correlation and the GNN approaches. **a**,**b**, The group mean SC was significantly correlated with mean FC in the HCP-YA (**a**, *r* = 0.396, *P* < 1×10^-16^) and HCP-D (**b**, *r* = 0.420, *P* < 1×10^-16^) datasets, based on the Pearson correlation. **c**,**d**, The GNN accurately predicted the group mean FC from the mean SC in both datasets. The predicted mean FC was significantly correlated with mean empirical FC in the HCP-YA (**c**, *r* = 0.953, *P* < 1×10^-16^) and HCP-D (**d**, *r* = 0.948, *P* < 1×10^-16^) datasets. **e**,**f**, At the individual participant level, the GNN model returned a much higher structure-function coupling than the correlation approach in both HCP-YA (**e**, correlation: *r* =0.297 ± 0.030; GNN: *r* =0.685 ± 0.046) and HCP-D (**f**, correlation: *r* =0.306 ± 0.028; GNN: 0.663 ± 0.047) datasets. Moreover, rewiring the SC network by disrupting the network topology and keeping the degree and strength distribution substantially reduced individuals’ structure-function coupling using GNNs in both HCP-YA (**e**, GNN: *r* = 0.308 ± 0.041) and HCP-D (**f**, GNN: *r* = 0.286 ± 0.029) datasets. This suggests that the structure-function coupling captured by GNNs was mainly driven by the network topology. **g**,**h**, The regional structure-function coupling by the correlation approach for HCP-YA (**g**) and HCP-D (**h**) datasets. **i**,**j**, The regional structure-function coupling by the GNN model for HCP-YA (**i**) and HCP-D (**j**) datasets. FC, functional connectivity; SC, structural connectivity; GNN, graph neural network.

It is noteworthy that correlation approaches only accounts for direct connections between nodes, while FC can emerge from polysynaptic communication on structural connectome (Avena-Koenigsberger et al., 2017; Seguin, Sporns, et al., 2023). A recent study indicated that a feed-forward fully-connected neural network accurately predicted FC from SC with higher accuracy than coupling with traditional biophysical models (Sarwar et al., 2021). Neudorf et al. further enhanced the prediction accuracy of functional connectivity by employing GNNs (Neudorf et al., 2022), which are designed for topological data (Zhou et al., 2020). Building on these studies, we proposed a GNN framework integrating graph convolutional network (GCN) and multi-layer perceptron (MLP) to predict FC from SC. At the group level, the Pearson correlation between the predicted and empirical FCs was *r* = 0.953 (*P* < 1×10^-16^; **Fig. 2c**) for the HCP-YA and *r* = 0.948 (*P* < 1×10^-16^; **Fig. 2d**) for the HCP-D dataset, suggesting a strong structure-function coupling captured by GNN models. At the individual level, GNN predicted FC from SC with an accuracy ranging from *r* = 0.514 to *r* = 0.782 (*r* = 0.685 ± 0.046; **Fig. 2e**) across HCP-YA participants and from *r* = 0.451 to *r* = 0.752 (*r* = 0.663 ± 0.047; **Fig. 2f**) across HCP-D participants. The lowest individual-level structure-function coupling from the GNN was higher than the strongest coupling from the correlation approach in both datasets. Overall, consistent with prior work from Neudorf et al. (2022), these results suggest that the GNN model captures a much stronger structure-function coupling than correlation approaches.

We further explored how network topology contributes to structure-function coupling prediction when using GNNs. To do this, we rewired the SC to randomize the network topology while maintaining the distribution of the nodal degree and strength (Maslov & Sneppen, 2002; Vasa & Misic, 2022). We trained GNN models using rewired SC in the training data to predict the FC in the test data. We found that GNN trained on rewired SC could only predict FC with an accuracy of *r* = 0.308 ± 0.041 (**Fig. 2e**) for individual participants in HCP-YA and *r* = 0.286 ± 0.029 (**Fig. 2f**) in HCP-D. Therefore, rewiring SC reduced participants’ average variance accounted for in the GNN-based structure-function coupling by 38% (empirical: mean *R^2^* = 0.471 ± 0.061; rewired: mean *R^2^* = 0.096 ± 0.027) in HCP-YA, and by 36% (empirical: mean *R^2^* = 0.441 ± 0.059; rewired: mean *R^2^* = 0.083 ± 0.017) in HCP-D. Overall, these results suggest that network topology drives GNN-based structure-function coupling.

After demonstrating topology driven structure-function coupling at the whole-brain level, we evaluated structure-function coupling at the regional level for both the linear correlation and GNN approaches. Specifically, the structural profile of each cortical region was defined as its connected SC edges for the correlation approach and the predicted FC of its connected edges for the GNN model. The functional profile of each cortical region was defined as the empirical FC of connected edges. Regional structure-function coupling was measured as the Pearson correlation between the structural and functional profiles for each cortical region. For both the linear correlation and the GNN approaches, regional structure-function coupling was heterogeneously distributed across the cortex, with a higher value in the primary sensorimotor cortices and a lower value in the higher-order association cortices, which was consistent with prior literature (Baum et al., 2020; Vazquez-Rodriguez et al., 2019). Particularly, the regional structure-function coupling ranged from 0.074 to 0.575 for HCP-YA (**Fig. 2g**) and from 0.065 to 0.640 for HCP-D (**Fig. 2h**) across all 400 cortical regions for the correlation approach. Using the GNN model, the regional structure-function coupling increased substantially, ranging from 0.326 to 0.809 for HCP-YA (**Fig. 2i**) and from 0.263 to 0.801 for HCP-D (**Fig. 2j**) across the cortical regions.

Previous studies have demonstrated that the cortical distribution of structure-function coupling aligns with functional hierarchy across the human cortex (Baum et al., 2020; Sydnor et al., 2021; Vazquez-Rodriguez et al., 2019). To replicate this finding, we acquired an a priori cortical map of the sensorimotor-association cortical axis (**Fig. S2a**), representing a unified cortical hierarchy along which various neurobiological properties are systematically distributed (Sydnor et al., 2021). The cortical regions were continuously ranked along this axis, with the primary sensorimotor cortices occupying the lowest ranks and higher-order association cortices occupying the highest ranks. We observed negative associations between structure-function coupling and sensorimotor-association axis ranks across all cortical regions using both the linear correlation (HCP-YA: Spearman’s rho = -0.54, *P*_spin_ < 0.001, **Fig. S2b**; HCP-D: Spearman’s rho = -0.50, *P*_spin_ < 0.001, **Fig. S2c**) and GNN (HCP-YA: Spearman’s rho = -0.48, *P*_spin_ < 0.001, **Fig. S2d**; HCP-D: Spearman’s rho = -0.29, *P*_spin_ = 0.003, **Fig. S2e**) approaches.

### 3.2. Structure-function coupling is primarily dominated by group-common effects, with subtle but significant individual-specific effects

Having found that structure-function coupling was reliable and driven by network topology, we next examined the magnitudes of group-common and individual-specific effects in structure-function coupling, utilizing both the correlation approach and the GNN model. To achieve this, we followed the approach used by Gratton et al. for analyzing group and individual effects of functional networks (Gratton et al., 2018) and constructed a participant-by-participant matrix of structure-function coupling for both models. The diagonal elements of the matrix quantify the structure-function coupling within individuals, whereas the off-diagonal elements quantify the coupling between one participant’s SC (or predicted FC when using the GNN) and another participant’s FC (**Fig. 3a**). We designated the within-individual coupling as “matched coupling” and the between-individual coupling as “mismatched coupling”.

**Fig. 3.**
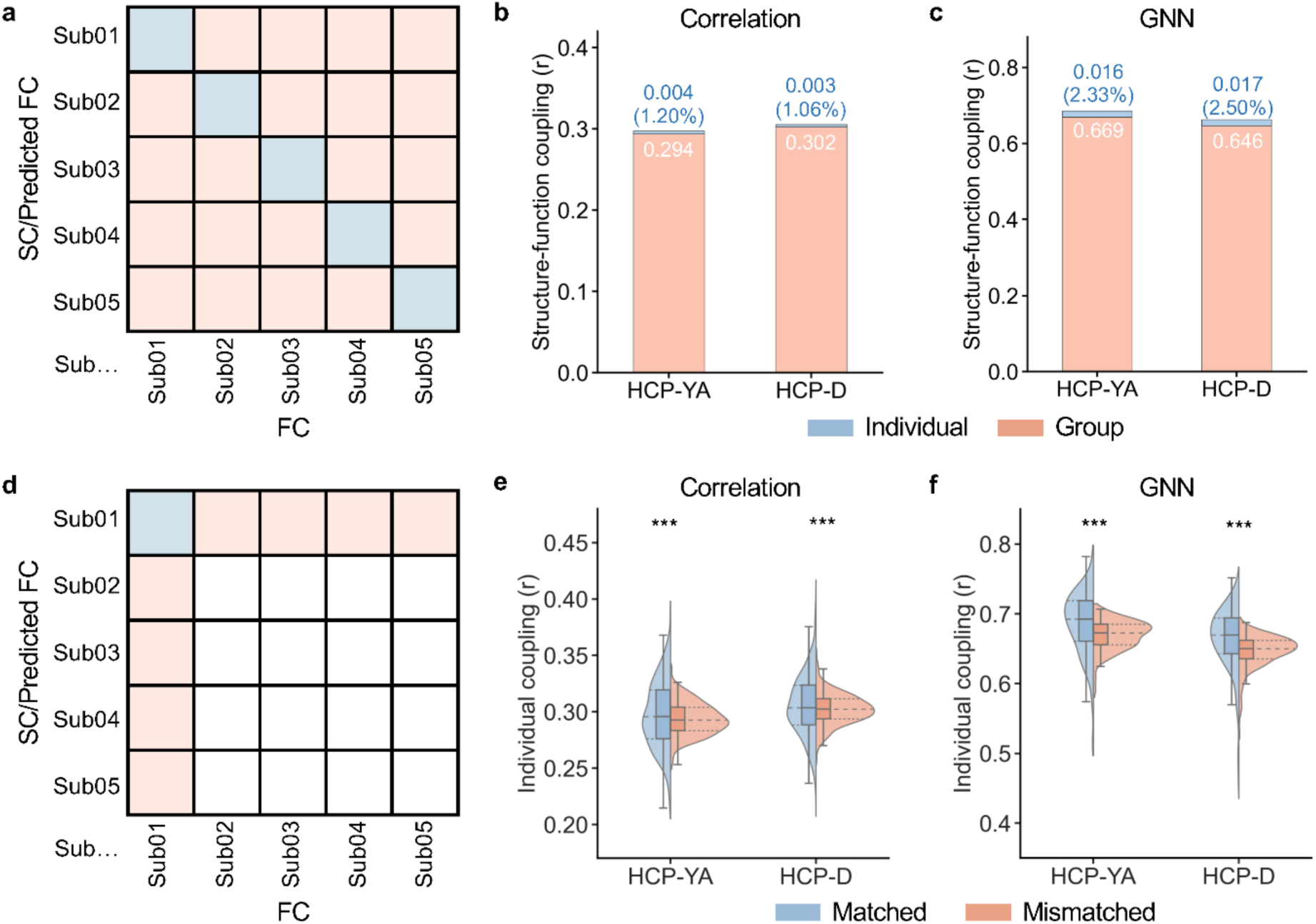
The group-common and individual-specific effects of structure-function coupling. **a**, The matrix of structure-function coupling between each pair of participants. Diagonal elements (cold color) represent the within-subject coupling (i.e., matched coupling), while off-diagonal elements (warm color) represent the coupling between one participant’s SC or predicted FC and another participant’s FC (i.e., mismatched coupling). SC was used for the correlation approach while the predicted FC was used for the GNN model to calculate the structure-function coupling. The total effects were defined as the average value of the diagonal elements; the group-common effects were defined as the average value of the off-diagonal elements; and the individual-specific effects were defined as their difference. **b**,**c**, The group and individual effects were estimated using the correlation approach (**b**, HCP-YA: 0.294/0.004 [group/individual]; HCP-D: 0.302/0.003) and the GNN model (**c**, HCP-YA: 0.669/0.016; HCP-D: 0.646/0.017). **d**, The matched coupling and the average of all mismatched coupling were extracted for each participant and then were statistically compared across participants using two-tailed paired t-tests. **e**, **f**, The matched individual coupling was significantly higher than the mismatched coupling for both the correlation approach (**e**) and the GNN model (**f**), suggesting statistically significant individual-specific effects of structure-function coupling. *** indicates *P* < 0.0001, two-tailed paired t-test. FC, functional connectivity; SC, structural connectivity; GNN, graph neural network.

As expected, the average of all pairs of matched coupling (correlation: [HCP-YA: *r* = 0.297; HCP-D: *r* = 0.306]; GNN: [HCP-YA: *r* = 0.685; HCP-D: *r* = 0.663]) was higher than the average of all pairs of mismatched couplings (correlation: [HCP-YA: *r* = 0.294; HCP-D: *r* = 0.302]; GNN: [HCP-YA: *r* = 0.669; HCP-D: *r* = 0.646]) for both the correlation approach and the GNN model. We defined the total effects of structure-function coupling as the average matched coupling, group-common effects as the average mismatched coupling, and individual-specific effects as the difference between the average matched coupling and average mismatched coupling (i.e., total effects minus group-common effects). Using the correlation approach, our results revealed that individual effects accounted for 1.20% of the total effects of structure-function coupling in HCP-YA and 1.06% in HCP-D (**Fig. 3b**). By contrast, the individual effects of the GNN model accounted for 2.33% and 2.50% of the total effects in HCP-YA and HCP-D, respectively (**Fig. 3c**). These results indicate that the GNN outperformed the correlation approach in capturing individual effects, suggesting that some individual effects of structure-function coupling can be explained by the nonlinear and higher-order network topology. Moreover, both models demonstrated that structure-function coupling was primarily dominated by group-common factors (more than 97%) rather than individual-specific traits (less than 3%).

As the individual effects of structure-function coupling were minor for both models, we next evaluated whether the individual effects were statistically significant. To test this, we extracted the matched coupling and the average of all mismatched coupling for each participant (**Fig. 3d**). We compared matched coupling and average mismatched coupling across all participants using two-tailed paired t-tests. The results indicated that the matched coupling was significantly higher than the mismatched coupling in both the correlation approach (HCP-YA: *P* < 0.0001; HCP-D: *P* < 0.0001; **Fig. 3e**) and the GNN model (HCP-YA: *P* < 0.0001; HCP-D: *P* < 0.0001; **Fig. 3f**). These findings confirm that the individual effects of structure-function coupling were statistically significant, as demonstrated by two independent datasets and two distinct approaches.

Overall, our findings suggest that individual structure-function coupling primarily reflects group-common characteristics, with subtle yet significant individual-specific effects.

### 3.3. Group and individual effects of structure-function coupling are organized on the cortex along the sensorimotor-association axis

Having identified the magnitudes of group-common and individual-specific effects in structure-function coupling at the whole-brain level, we evaluated how group and individual effects were distributed across the cortex at the regional level. We hypothesized that the cortical distributions of both group and individual effects would align with the sensorimotor-association cortical axis (**Fig. 4a**).

**Fig. 4.**
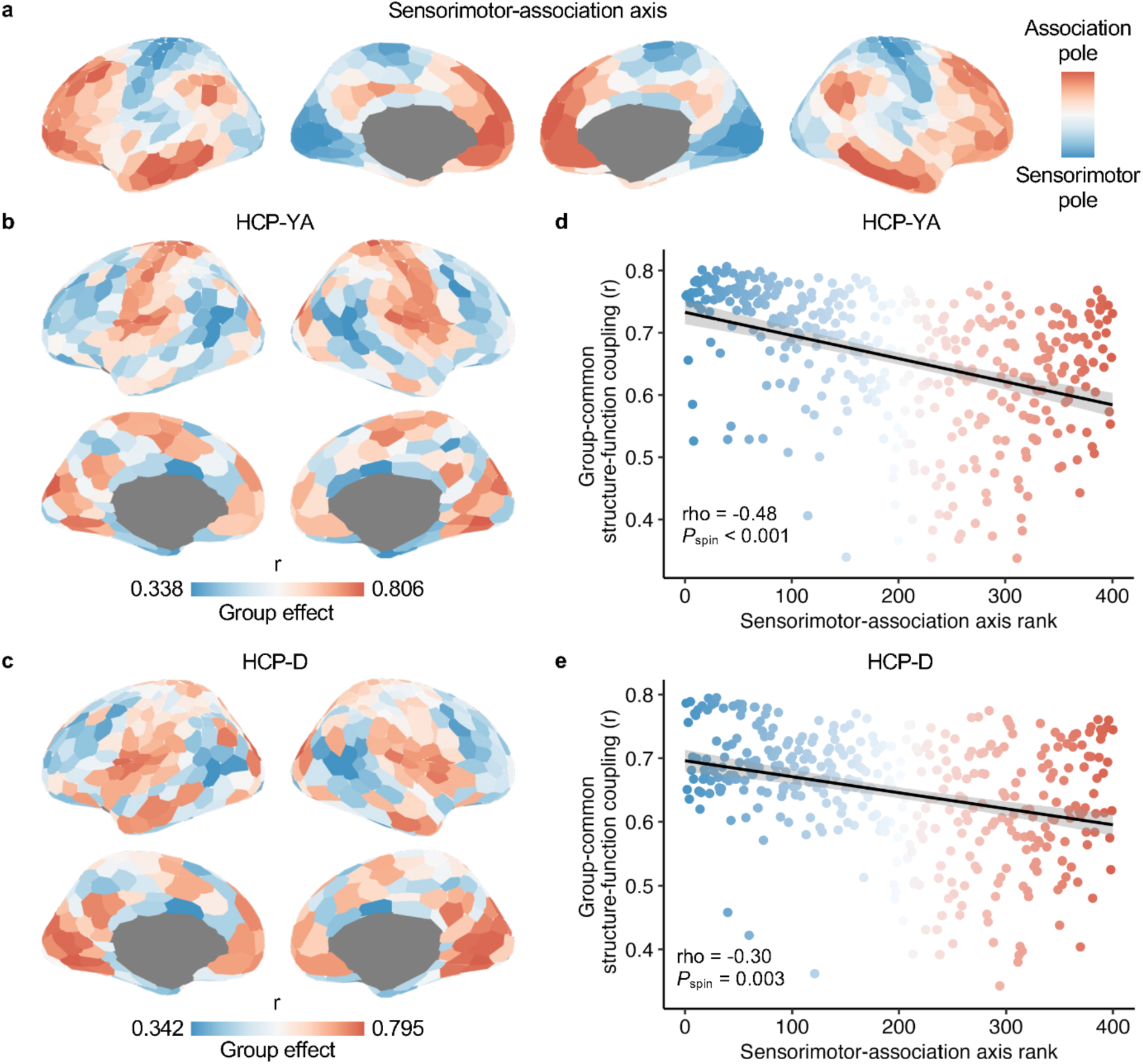
GNN-derived regional group-common structure-function effects inversely align with the sensorimotor-association cortical axis. **a**, The cortical map of the sensorimotor-association axis was derived from Sydnor et al. (Sydnor et al., 2021), where warm colors represent association cortices with higher ranks and cold colors represent sensorimotor cortices with lower ranks. **b**,**c**, The cortical map of group-common effects in regional structure-function coupling estimated by the GNN model in the HCP-YA (**b**) and HCP-D (**c**) datasets. **d**,**e**, The regional group effects of structure-function coupling were negatively correlated with the sensorimotor-association axis ranks in the HCP-YA (**d**, Spearman’s rho = -0.48, *P*_spin_ < 0.001) and HCP-D (**e**, Spearman’s rho = -0.30, *P*_spin_ = 0.003) datasets. The outliers (mean ± 3×SD) were excluded from the Spearman’s rank correlation analysis. Each point in the scatter plot represents a cortical region and is colored by its rank in the sensorimotor-association axis. GNN, graph neural network.

We calculated the group and individual effects of the GNN-based structure-function coupling at the regional level. We constructed a participant-by-participant matrix of the GNN-based coupling and averaged all mismatched couplings for each cortical region, resulting in regional group effects. We observed that the group effects were higher in the primary sensorimotor cortices and lower in the higher-order association cortices in both the HCP-YA (**Fig. 4b**) and HCP-D (**Fig. 4c**) datasets. Using the Spearman’s rank correlation and a conservative spin-based spatial permutation test (Alexander-Bloch et al., 2018), we found that the cortical distributions of the regional group effects were highly reproducible between the two datasets (Spearman’s rho = 0.86, *P*_spin_ < 0.001). Using the correlation approach to evaluate regional structure-function coupling, we observed that the cortical distribution of group effects was similar to that using the GNN model in both the HCP-YA and HCP-D datasets (**Fig. S3a**).

Next, we quantitatively evaluated whether the group effects of the regional structure-function coupling were organized along the sensorimotor-association cortical axis. Using the Spearman’s rank correlation, we found that the group effects, based on the GNN model, were negatively correlated with the ranks of the sensorimotor-association axis across all cortical regions in both the HCP-YA (Spearman’s rho = -0.48, *P*_spin_ < 0.001; **Fig. 4d**) and HCP-D (Spearman’s rho = -0.30, *P*_spin_ = 0.003; **Fig. 4e**) datasets. The sensorimotor pole of the cortical axis showed a higher group effect, and the association pole showed a lower group effect. Additionally, using a correlation approach to evaluate structure-function coupling, we consistently observed a negative association between regional group effects and sensorimotor-association axis ranks in both HCP-YA (Spearman’s rho = -0.54, *P*_spin_ < 0.001; **Fig. S3b**) and HCP-D (Spearman’s rho = -0.50, *P*_spin_ < 0.001; **Fig. S3c**). These results suggest that the group effects of structure-function coupling were spatially patterned on the cortex along the sensorimotor-association axis.

Next, we examined the cortical distribution of the individual-specific effects of regional structure-function coupling. Using regional participant-by-participant matrices of GNN-based structure-function coupling, we calculated individual effects as the difference between the average matched and mismatched coupling for each cortical region. We observed that individual-specific effects were weaker in the sensorimotor cortices and stronger in the association cortices in both the HCP-YA (**Fig. 5a**) and HCP-D (**Fig. 5b**) datasets. The cortical distribution of individual effects was significantly aligned between the two datasets (Spearman’s rho = 0.48, *P*_spin_ < 0.001). Using the correlation-based structure-function coupling, we observed a similar cortical pattern of individual effects with those using the GNN model for both datasets (**Fig. S3d**).

**Fig. 5.**
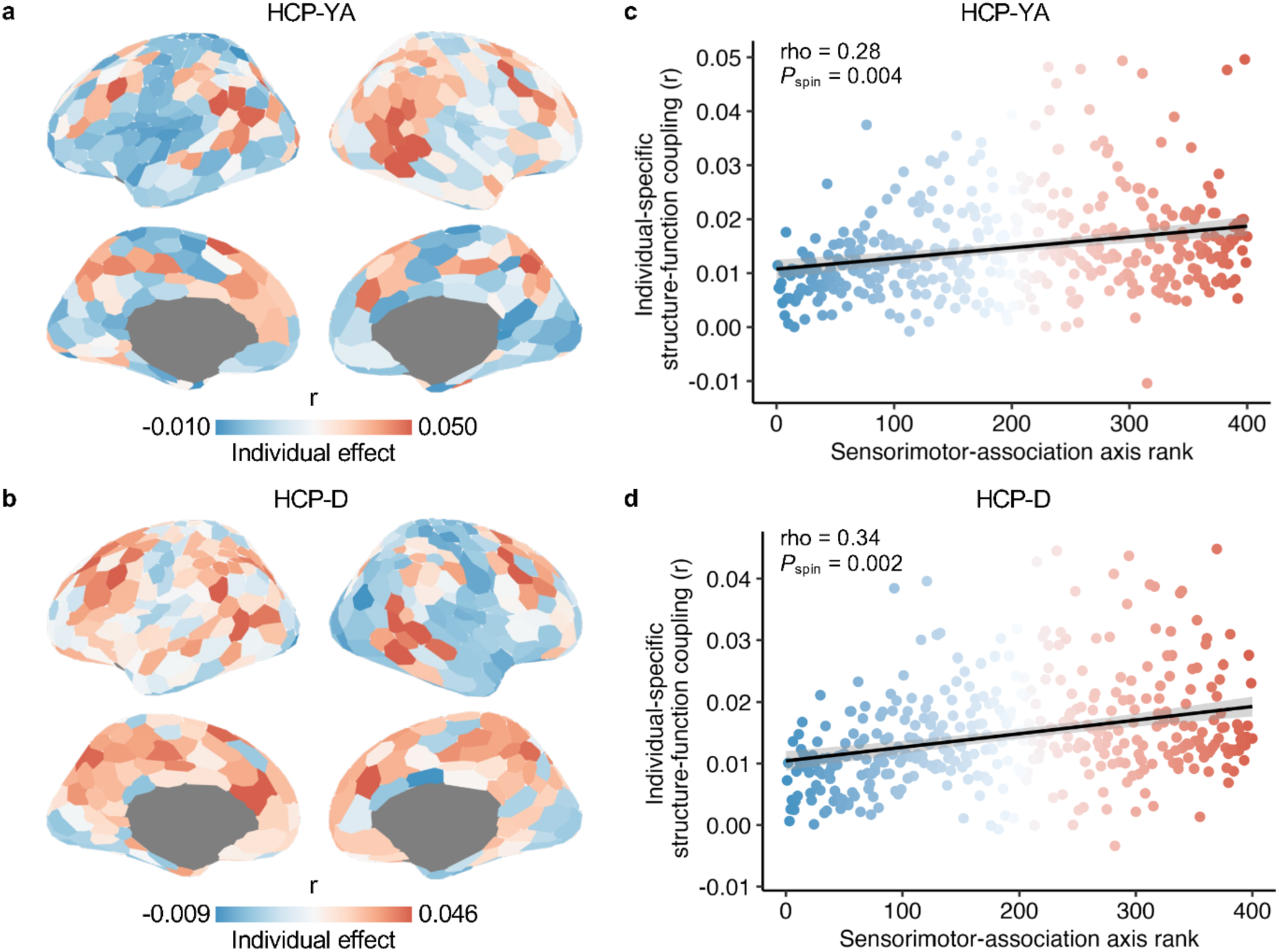
GNN-derived regional individual-specific effects of structure-function positively align with the sensorimotor-association cortical axis. **a**,**b**, The cortical map of individual-specific effects in structure-function coupling as estimated by the GNN model in the HCP-YA (**a**) and HCP-D (**b**) datasets. **c**,**d**, The individual-specific effects were positively correlated with the ranks in the sensorimotor-association axis across all cortical regions in the HCP-YA (**c**, Spearman’s rho = 0.28, *P*_spin_ = 0.004) and HCP-D (**d**, Spearman’s rho = 0.34, *P*_spin_ = 0.002) datasets. The outliers (mean ± 3×SD) were excluded from the Spearman’s rank correlation analysis. Each point in the scatter plot represents a cortical region and is colored by its cortical axis rank. GNN, graph neural network.

By quantifying the alignment with the sensorimotor-association cortical axis, we found a significant positive correlation between the individual effects and the ranks of this cortical axis across all cortical regions in both the HCP-YA (Spearman’s rho = 0.28, *P*_spin_ = 0.004; **Fig. 5c**) and HCP-D (Spearman’s rho = 0.34, *P*_spin_ = 0.002; **Fig. 5d**) datasets. These results indicate an opposite cortical pattern in individual-specific effects compared to group-common effects. The sensorimotor pole of the axis had a lower individual effect, and the association pole showed a higher individual effect. However, using the correlation approach, the association between regional individual effects and cortical axis ranks was not significant in either the HCP-YA (Spearman’s rho = -0.002, *P*_spin_ = 0.510; **Fig. S3e**) or HCP-D (Spearman’s rho = -0.15, *P*_spin_ = 0.127; **Fig. S3f**) datasets. These results suggest that GNN is better at capturing individual-specific effects of structure-function coupling in the human brain.

### 3.4. Sensitivity analyses

We performed three additional analyses to ensure that our results were robust to the methodological choices. First, to ensure that our findings were robust to the partition of the training and test sets, we conducted repeated 2-fold cross-validations, randomly splitting the dataset into two halves 100 times. The average group-level structure-function coupling across all folds using GNNs was *r* = 0.948 ± 0.006 in the HCP-YA and *r* = 0.946 *±* 0.004 in the HCP-D. The average of the participants’ mean individual-level coupling across all folds was *r* = 0.685 ± 0.005 for the HCP-YA participants and *r* = 0.657 ± 0.002 for the HCP-D participants. The individual effects accounted for 2.30% ± 0.13% in HCP-YA and 2.62% ± 0.07% in HCP-D, on average across splits, with all values being significant (*P* < 0.001 for all splits using the paired t-test). Moreover, the average Spearman’s rank correlation between group effects and sensorimotor-association ranks was rho = -0.446 ± 0.022 in the HCP-YA (*P*_spin_ < 0.001 for all splits) and rho = -0.273 ± 0.013 in the HCP-D (all *P*_spin_ < 0.001). The average correlation between individual effects and sensorimotor-association ranks was rho = 0.272 ± 0.059 in the HCP-YA (all *P*_spin_ < 0.05) and rho = 0.255 ± 0.038 in the HCP-D (all *P*_spin_ < 0.05). These results demonstrated that the GNN’s performance is robust to the variation of train-test splits.

In this study, we defined the individual effects of structure-function coupling by subtracting the average mismatched coupling (between-subject coupling) from the matched coupling (within-subject coupling). This measurement may be influenced by the magnitude of the total effects, as larger total effects can potentially generate larger individual effects. To mitigate the potential scaling issue, we defined the normalized individual effects as the proportion of the individual effect to the total effect and re-evaluated the alignment between the normalized individual effects and the sensorimotor-association cortical axis ranks. Similar to our main results, we found that the normalized individual effects were lower in the sensorimotor cortices and higher in the association cortices (**Fig. S4a** & **S4b**). Moreover, we observed a significant positive correlation between the normalized individual effects and cortical axis ranks in the HCP-YA (Spearman’s rho = 0.33, *P*_spin_ < 0.001; **Fig. S4c**) and HCP-D (Spearman’s rho = 0.36, *P*_spin_ = 0.001; **Fig. S4d**) datasets.

Finally, we validated our findings of GNN-based structure-function coupling using another Schaefer-200 parcellation (Schaefer et al., 2018). We first confirmed the significance of individual-specific structure-function coupling effects. We observed significant individual effects of coupling with the Schaefer-200 (**Fig. S5a**). Furthermore, we demonstrated a negative association between group effects and the sensorimotor-association cortical axis ranks (**Fig. S5 b-d**), and a positive association between individual effects and cortical ranks (**Fig. S5 e-g**) across all regions. Overall, our main results are robust to the choice of cortical parcellation.

## 4. DISCUSSION

In this study, we used a graph neural network (GNN) framework to evaluate the magnitude of the group-common and individual-specific effects of structure-function coupling. As in prior work (Neudorf et al., 2022), we found that SC accurately predicted the FC of unseen individuals using the GNN model, and extended this work by demonstrating the prediction was driven by network topology. We observed that structure-function coupling was dominated by group-common characteristics; simultaneously, minor individual-specific effects were also noted to be significant. Finally, we found that both regional group and individual effects were hierarchically patterned across the cortex along the fundamental sensorimotor-association cortical axis. The sensorimotor pole of the axis showed a higher group effect and lower individual effect, whereas the association pole showed a higher individual effect and lower group effect in structure-function coupling. These results were consistent for the two independent high-quality datasets. Our findings emphasize the importance of considering group and individual effects in individual difference studies of structure-function coupling.

Understanding the dynamic communication process in the structural connectome is a central goal in neuroscience (Avena-Koenigsberger et al., 2017; Seguin, Sporns, et al., 2023). Prior studies have demonstrated that the SC of macroscale white matter tracts is associated with FC, as measured by the correlation between pairs of regional time series (Baum et al., 2020; Betzel et al., 2019; Deco et al., 2013; Demirtas et al., 2019; Gu et al., 2021; Hermundstad et al., 2013; Honey et al., 2009; Medaglia et al., 2018; Misic et al., 2016; Misic et al., 2015; Sarwar et al., 2021; Seguin, Jedynak, et al., 2023; Suárez et al., 2021; Valk et al., 2022; Vazquez-Rodriguez et al., 2019; Zamani Esfahlani et al., 2022). The cortical distribution of regional structure-function coupling aligns with the axis of the fundamental sensorimotor-association cortical hierarchy (Baum et al., 2020; Sydnor et al., 2021; Vazquez-Rodriguez et al., 2019). Moreover, structure-function coupling changes with age and is related to individual differences in cognition and psychopathology (Baum et al., 2020; Jiang et al., 2020; Jiang et al., 2019; Medaglia et al., 2018; Zamani Esfahlani et al., 2022). Building on these studies, our work provides a systematic examination of the group-common and individual-specific effects of the structure-function coupling at the global and regional levels using both the correlation approach and advanced GNN.

Using the GNN framework, we found that SC reconstructed using diffusion MRI accurately predicted functional connectivity in unseen individuals using resting-state fMRI. In line with our results, previous studies have consistently demonstrated a robust coupling between SC and FC using a variety of interpretable models, such as correlation approaches (Baum et al., 2020; Gu et al., 2021; Honey et al., 2009; Vazquez-Rodriguez et al., 2019), biophysical models (Deco et al., 2013), communication models (Betzel et al., 2019; Seguin, Jedynak, et al., 2023; Seguin et al., 2022; Seguin, Sporns, et al., 2023; Zamani Esfahlani et al., 2022), Riemannian approaches (Benkarim et al., 2022; Deslauriers-Gauthier et al., 2022). However, these interpretable models typically return relatively small structure-function coupling values, even when accounting for indirect communication, suggesting a potentially imperfect alignment between structural and functional connectivity (Seguin, Sporns, et al., 2023; Suárez et al., 2020; Valk et al., 2022). Recently, Sarwar et al. found that a fully-connected neural network predicted FC with much higher accuracy than interpretable models (Sarwar et al., 2021), suggesting that structure-function coupling could be more significant than previously imagined. Two recent studies employed GNNs to investigate structure-function coupling; however, they did not evaluate how network topology contributed to the coupling prediction, despite utilizing network topological information being a core feature of GNNs (Hong et al., 2023; Neudorf et al., 2022). Our results extended these findings by demonstrating that the GNN’s performance was primarily attributed to network topology. Specifically, by randomizing the network topology of SC, our GNN model reduced the variance accounted by more than 36%. These results are also consistent with prior research showing that GNNs are effective at capturing topological representations of network data, such as cellular networks (Wu et al., 2022) and protein networks (Gao et al., 2023).

Our results also demonstrated that structure-function coupling was dominated by group-common effects, but the remaining minor individual effects were also significant. Robust group effects could underlie past success in the estimation of structure-function coupling at the group level (Deco et al., 2013; Demirtas et al., 2019; Honey et al., 2009; Misic et al., 2016; Misic et al., 2015; Suárez et al., 2021; Vazquez-Rodriguez et al., 2019). This result aligns with previous studies showing that functional networks are largely determined by the group-common organizational principle (Gratton et al., 2018) and that structural networks are even less variable than functional networks (Zimmermann et al., 2019). Directly supporting our results, a recent study observed that structure-function coupling, based on a correlation approach, was dominated by group effects, consistent across six datasets with different acquisitions or processing methods (Zimmermann et al., 2019). However, individual effects of structure-function coupling were only observed in one of the six datasets (Zimmermann et al., 2019). Here, using an advanced GNN approach and two independent, high-quality datasets, we demonstrated that the individual effects of structure-function coupling were also robust and reproducible. Moreover, our results indicated that the individual effects with the GNN approach were much larger than those with the correlation approach, which potentially explained the nonstable individual effects in a previous study using Pearson correlation (Zimmermann et al., 2019). Recently, Smolder et al. questioned whether structural connectivity predicts functional connectivity at the individual level (Smolders et al., 2024). Our findings provide a reliable and quantitative demonstration of the structure-function coupling at the individual level. Consistently, a recent study also reported the significant presence of individual-specific effects of structure-function coupling using fully-connected neural networks (Zalesky et al., 2024). Future research is needed to explore methods for improving interpretable network communication models to better detect individual effects of structure-function coupling.

The sensorimotor-association cortical hierarchy has been identified as a unifying cortical organizing principle for diverse neurobiological properties, including structure, function, metabolism, transcription, and evolution (Baum et al., 2020; Burt et al., 2018; Demirtas et al., 2019; Hill et al., 2010; Huntenburg et al., 2018; Margulies et al., 2016; Murray et al., 2014; Raut et al., 2020; Sydnor et al., 2021; Vazquez-Rodriguez et al., 2019). Recent studies have demonstrated that the cortical distribution of structure-function coupling aligns with the sensorimotor-association axis (Baum et al., 2020; Vazquez-Rodriguez et al., 2019). Consistent with these previous reports, we found that both the group and individual effects of regional structure-function coupling were hierarchically distributed across the cortex along the sensorimotor-association cortical axis. More specifically, we observed that the higher-order association cortex, which is located at the top end of the cortical axis, displayed the highest individual effects and lowest group effects, whereas the primary sensorimotor cortex displayed the highest group effects and lowest individual effects. Both structural and functional connectivity exhibit the highest inter-individual variability in the association cortex and the lowest in the sensorimotor cortex (Huang et al., 2023; Mueller et al., 2013), which partly supports our findings. This greater individual variability in the association cortex may result in more pronounced individual-specific coupling effects, whereas the lower variability in the sensorimotor cortex may lead to reduced individual-specific coupling effects. Moreover, the sensorimotor cortex primarily comprises circuits with relatively simple and canonical feedforward and feedback connectivity patterns (Felleman & Van Essen, 1991), which could potentially result in high cross-individual similarity in structure-function coupling. In contrast, the association cortex exhibits a non-canonical circuit architecture with complicated and distributed connections (Goldman-Rakic, 1988), which could lead to individual diversity in structure-function coupling.

This study has several potential limitations. First, precisely reconstructing individuals’ white matter SC is challenging because of the inherent limitations of diffusion MRI-based fiber tractography. In this study, we used state-of-the-art probabilistic fiber tractography with multi-shell, multi-tissue constrained spherical deconvolution (Jeurissen et al., 2014) and applied anatomically constrained tractography (Smith et al., 2012) and spherical deconvolution-informed filtering of tractograms (Smith et al., 2015) to improve biological accuracy. Moreover, consistency-based thresholding was used to reduce the influence of false-positive connections (Baum et al., 2020). Second, the current study did not analyze the structure-function coupling of the subcortical and cerebellar structures. Future studies could explore both the group-common and individual-specific effects of structure-function coupling in these areas. Finally, although the GNN accurately predicted FC with structural network topology, its black-box nature prevented us from understanding the underlying communication mechanism supporting this prediction. However, this result provides a benchmark for optimizing interpretable models to explicitly explain how the structural connectome supports the functional connectivity.

Notwithstanding these limitations, we reliably demonstrated that the structure-function coupling is dominated by the group-common effects and the minor yet significant individual-specific effects using advanced GNN models. The individual coupling effects were the highest in the association cortex, which is related to the prolonged development of higher-order cognition and confers diverse psychopathologies (Sydnor et al., 2021). This implies that structure-function coupling could be a potential neuromarker for tracking individual differences in cognitive development and vulnerability to mental disorders. However, the minor individual effects also suggest caution in interpreting the individual differences in structure-function coupling among individuals. As structural connectivity pathways facilitate the propagation of neurostimulation-induced activity (Seguin, Jedynak, et al., 2023; Sydnor et al., 2022), our findings may have implications for the clinical practice of neurostimulation. The dominant group effects in structure-function coupling could explain why the same neurostimulation target (i.e., dorsal lateral prefrontal cortex) has benefited many different patients (Hyde et al., 2022). Simultaneously, significant individual-specific effects could explain why personalized stimulation targets have improved intervention effects for certain patients (Klooster et al., 2022).

## DATA AND CODE AVAILABILITY

The HCP-YA and HCP-D datasets are available at https://db.humanconnectome.org/. All code used to perform the analyses in this study can be found at https://github.com/CuiLabCIBR/GNN_SC_FC.

## AUTHOR CONTRIBUTIONS

Z.C. and P.C. designed the study. P.C. and H.Y. performed the analyses with support from T.S.O. H.Y. and X.X. completed data preprocessing. X.Z., H.J. and J.H. replicated the model training and testing. H.Y and P.C. created the figures. Z.C. and T.S.O. supervised the project. R.C. provided resources. P.C., H.Y., and Z.C. wrote the manuscript with review and editing from all other authors.

## FUNDING

This work is supported by the STI 2030-Major Projects (2022ZD0211300), Beijing Nova Program (Z211100002121002), and CIBR funds. Data were provided [in part] by the Human Connectome Project, WU-Minn Consortium (Principal Investigators: David Van Essen and Kamil Ugurbil; 1U54MH091657) funded by the 16 NIH Institutes and Centers that support the NIH Blueprint for Neuroscience Research; and by the McDonnell Center for Systems Neuroscience at Washington University. The HCP-Development 2.0 Release data used in this report came from DOI: 10.15154/1520708. Research reported in this publication was supported by the National Institute of Mental Health of the National Institutes of Health under Award Number U01MH109589 and by funds provided by the McDonnell Center for Systems Neuroscience at Washington University in St. Louis.

## DECLARATION OF COMPETING INTERESTS

The authors declare no competing interests.

## Notes

### Competing Interest Statement

The authors have declared no competing interest.

### Summary of Updates

Add extra analyses and improve the writing.

## REFERENCES

Alexander-Bloch, A. F., Shou, H., Liu, S., Satterthwaite, T. D., Glahn, D. C., Shinohara, R. T., Vandekar, S. N., & Raznahan, A. (2018). On testing for spatial correspondence between maps of human brain structure and function. Neuroimage, 178, 540–551. 10.1016/j.neuroimage.2018.05.070

Andersson, J. L. R., Graham, M. S., Zsoldos, E., & Sotiropoulos, S. N. (2016). Incorporating outlier detection and replacement into a non-parametric framework for movement and distortion correction of diffusion MR images. Neuroimage, 141, 556–572. 10.1016/j.neuroimage.2016.06.058

Avena-Koenigsberger, A., Misic, B., & Sporns, O. (2017). Communication dynamics in complex brain networks. Nat Rev Neurosci., 19(1), 17–33. 10.1038/nrn.2017.149

Bassett, D. S., & Bullmore, E. (2006). Small-world brain networks. Neuroscientist, 12(6), 512–523. 10.1177/1073858406293182

Baum, G. L., Cui, Z., Roalf, D. R., Ciric, R., Betzel, R. F., Larsen, B., Cieslak, M., Cook, P. A., Xia, C. H., Moore, T. M., Ruparel, K., Oathes, D. J., Alexander-Bloch, A. F., Shinohara, R. T., Raznahan, A., Gur, R. E., Gur, R. C., Bassett, D. S., & Satterthwaite, T. D. (2020). Development of structure-function coupling in human brain networks during youth. Proc Natl Acad Sci U S A, 117(1), 771–778. 10.1073/pnas.1912034117

Benkarim, O., Paquola, C., Park, B.-y., Royer, J., Rodríguez-Cruces, R., Vos de Wael, R., Misic, B., Piella, G., & Bernhardt, B. C. (2022). A Riemannian approach to predicting brain function from the structural connectome. Neuroimage, 257, 119299. 10.1016/j.neuroimage.2022.119299

Betzel, R. F., Medaglia, J. D., Kahn, A. E., Soffer, J., Schonhaut, D. R., & Bassett, D. S. (2019). Structural, geometric and genetic factors predict interregional brain connectivity patterns probed by electrocorticography. Nat Biomed Eng, 3(11), 902–916. 10.1038/s41551-019-0404-5

Burt, J. B., Demirtas, M., Eckner, W. J., Navejar, N. M., Ji, J. L., Martin, W. J., Bernacchia, A., Anticevic, A., & Murray, J. D. (2018). Hierarchy of transcriptomic specialization across human cortex captured by structural neuroimaging topography. Nat Neurosci, 21(9), 1251–1259. 10.1038/s41593-018-0195-0

Cieslak, M., Cook, P. A., He, X., Yeh, F.-C., Dhollander, T., Adebimpe, A., Aguirre, G. K., Bassett, D. S., Betzel, R. F., Bourque, J., Cabral, L. M., Davatzikos, C., Detre, J. A., Earl, E., Elliott, M. A., Fadnavis, S., Fair, D. A., Foran, W., Fotiadis, P., … Satterthwaite, T. D. (2021). QSIPrep: an integrative platform for preprocessing and reconstructing diffusion MRI data. Nature Methods, 18(7), 775–778. 10.1038/s41592-021-01185-5

Ciric, R., Wolf, D. H., Power, J. D., Roalf, D. R., Baum, G. L., Ruparel, K., Shinohara, R. T., Elliott, M. A., Eickhoff, S. B., Davatzikos, C., Gur, R. C., Gur, R. E., Bassett, D. S., & Satterthwaite, T. D. (2017). Benchmarking of participant-level confound regression strategies for the control of motion artifact in studies of functional connectivity. Neuroimage, 154, 174–187. 10.1016/j.neuroimage.2017.03.020

Deco, G., Ponce-Alvarez, A., Mantini, D., Romani, G. L., Hagmann, P., & Corbetta, M. (2013). Resting-state functional connectivity emerges from structurally and dynamically shaped slow linear fluctuations. J Neurosci, 33(27), 11239–11252. 10.1523/JNEUROSCI.1091-13.2013

Demirtas, M., Burt, J. B., Helmer, M., Ji, J. L., Adkinson, B. D., Glasser, M. F., Van Essen, D. C., Sotiropoulos, S. N., Anticevic, A., & Murray, J. D. (2019). Hierarchical Heterogeneity across Human Cortex Shapes Large-Scale Neural Dynamics. Neuron, 101(6), 1181–1194 e1113. 10.1016/j.neuron.2019.01.017

Deslauriers-Gauthier, S., Zucchelli, M., Laghrissi, H., & Deriche, R. (2022). A Riemannian Revisiting of Structure–Function Mapping Based on Eigenmodes [Original Research]. Frontiers in Neuroimaging, 1. https://www.frontiersin.org/journals/neuroimaging/articles/10.3389/fnimg.2022.85 0266

Faskowitz, J., Esfahlani, F. Z., Jo, Y., Sporns, O., & Betzel, R. F. (2020). Edge-centric functional network representations of human cerebral cortex reveal overlapping system-level architecture. Nature Neuroscience, 23(12), 1644–1654. 10.1038/s41593-020-00719-y

Felleman, D. J., & Van Essen, D. C. (1991). Distributed hierarchical processing in the primate cerebral cortex. Cereb Cortex, 1(1), 1–47. 10.1093/cercor/1.1.1-a

Fischl, B. (2012). FreeSurfer. Neuroimage, 62(2), 774–781.

Gao, Z., Jiang, C., Zhang, J., Jiang, X., Li, L., Zhao, P., Yang, H., Huang, Y., & Li, J. (2023). Hierarchical graph learning for protein-protein interaction. Nat Commun, 14(1), 1093. 10.1038/s41467-023-36736-1

Ge, J., Yang, G., Han, M., Zhou, S., Men, W., Qin, L., Lyu, B., Li, H., Wang, H., Rao, H., Cui, Z., Liu, H., Zuo, X. N., & Gao, J. H. (2023). Increasing diversity in connectomics with the Chinese Human Connectome Project. Nat Neurosci, 26(1), 163–172. 10.1038/s41593-022-01215-1

Glasser, M. F., Sotiropoulos, S. N., Wilson, J. A., Coalson, T. S., Fischl, B., Andersson, J. L., Xu, J., Jbabdi, S., Webster, M., Polimeni, J. R., Van Essen, D. C., & Jenkinson, M. (2013). The Minimal Preprocessing Pipelines for the Human Connectome Project. Neuroimage, 80, 105–124. 10.1016/j.neuroimage.2013.04.127

Goldman-Rakic, P. S. (1988). Topography of cognition: parallel distributed networks in primate association cortex. Annu Rev Neurosci, 11, 137–156. 10.1146/annurev.ne.11.030188.001033

Gratton, C., Laumann, T. O., Nielsen, A. N., Greene, D. J., Gordon, E. M., Gilmore, A. W., Nelson, S. M., Coalson, R. S., Snyder, A. Z., Schlaggar, B. L., Dosenbach, N. U. F., & Petersen, S. E. (2018). Functional Brain Networks Are Dominated by Stable Group and Individual Factors, Not Cognitive or Daily Variation. Neuron, 98(2), 439–452 e435. 10.1016/j.neuron.2018.03.035

Gu, Z., Jamison, K. W., Sabuncu, M. R., & Kuceyeski, A. (2021). Heritability and interindividual variability of regional structure-function coupling. Nat Commun, 12(1), 4894. 10.1038/s41467-021-25184-4

Hagmann, P., Cammoun, L., Gigandet, X., Meuli, R., Honey, C. J., Wedeen, V. J., & Sporns, O. (2008). Mapping the Structural Core of Human Cerebral Cortex. PLOS Biology, 6(7), e159. 10.1371/journal.pbio.0060159

Hermundstad, A. M., Bassett, D. S., Brown, K. S., Aminoff, E. M., Clewett, D., Freeman, S., Frithsen, A., Johnson, A., Tipper, C. M., Miller, M. B., Grafton, S. T., & Carlson, J. M. (2013). Structural foundations of resting-state and task-based functional connectivity in the human brain. Proc Natl Acad Sci U S A, 110(15), 6169–6174. 10.1073/pnas.1219562110

Hill, J., Inder, T., Neil, J., Dierker, D., Harwell, J., & Van Essen, D. (2010). Similar patterns of cortical expansion during human development and evolution. Proc Natl Acad Sci U S A, 107(29), 13135–13140. 10.1073/pnas.1001229107

Honey, C. J., Sporns, O., Cammoun, L., Gigandet, X., Thiran, J. P., Meuli, R., & Hagmann, P. (2009). Predicting human resting-state functional connectivity from structural connectivity. Proc Natl Acad Sci U S A, 106(6), 2035–2040. 10.1073/pnas.0811168106

Hong, Y., Cornea, E., Girault, J. B., Bagonis, M., Foster, M., Kim, S. H., Prieto, J. C., Chen, H., Gao, W., Styner, M. A., & Gilmore, J. H. (2023). Structural and functional connectome relationships in early childhood. Developmental Cognitive Neuroscience, 64, 101314. 10.1016/j.dcn.2023.101314

Huang, W., Chen, H., Liu, Z., Dong, X., Feng, G., Liu, G., Ma, G., Zhang, Z., Su, L., & Shu, N. (2023). Individual Variability in the Structural Connectivity Architecture of the Human Brain. bioRxiv, 2023.2001. 2011.523683. 10.1101/2023.01.11.523683

Huntenburg, J. M., Bazin, P. L., & Margulies, D. S. (2018). Large-Scale Gradients in Human Cortical Organization. Trends Cogn Sci, 22(1), 21–31. 10.1016/j.tics.2017.11.002

Hyde, J., Carr, H., Kelley, N., Seneviratne, R., Reed, C., Parlatini, V., Garner, M., Solmi, M., Rosson, S., Cortese, S., & Brandt, V. (2022). Efficacy of neurostimulation across mental disorders: systematic review and meta-analysis of 208 randomized controlled trials. Mol Psychiatry, 27(6), 2709–2719. 10.1038/s41380-022-01524-8

Jenkinson, M., Bannister, P., Brady, M., & Smith, S. (2002). Improved Optimization for the Robust and Accurate Linear Registration and Motion Correction of Brain Images. Neuroimage, 17(2), 825–841. 10.1006/nimg.2002.1132

Jeurissen, B., Tournier, J.-D., Dhollander, T., Connelly, A., & Sijbers, J. (2014). Multi-tissue constrained spherical deconvolution for improved analysis of multi-shell diffusion MRI data. Neuroimage, 103, 411–426. 10.1016/j.neuroimage.2014.07.061

Jiang, H., Zhu, R., Tian, S., Wang, H., Chen, Z., Wang, X., Shao, J., Qin, J., Shi, J., Liu, H., Chen, Y., Yao, Z., & Lu, Q. (2020). Structural-functional decoupling predicts suicide attempts in bipolar disorder patients with a current major depressive episode. Neuropsychopharmacology, 45(10), 1735–1742. 10.1038/s41386-020-0753-5

Jiang, X., Shen, Y., Yao, J., Zhang, L., Xu, L., Feng, R., Cai, L., Liu, J., Chen, W., & Wang, J. (2019). Connectome analysis of functional and structural hemispheric brain networks in major depressive disorder. Transl Psychiatry, 9(1), 136. 10.1038/s41398-019-0467-9

Kellner, E., Dhital, B., Kiselev, V. G., & Reisert, M. (2016). Gibbs-ringing artifact removal based on local subvoxel-shifts. Magnetic Resonance in Medicine, 76(5), 1574–1581. 10.1002/mrm.26054

Kingma, D. P., & Ba, J. (2017). Adam: A Method for Stochastic Optimization. 10.48550/arXiv.1412.6980

Kipf, T. N., & Welling, M. (2017). Semi-Supervised Classification with Graph Convolutional Networks. 10.48550/arXiv.1609.02907

Klooster, D. C. W., Ferguson, M. A., Boon, P., & Baeken, C. (2022). Personalizing Repetitive Transcranial Magnetic Stimulation Parameters for Depression Treatment Using Multimodal Neuroimaging. Biol Psychiatry Cogn Neurosci Neuroimaging, 7(6), 536–545. 10.1016/j.bpsc.2021.11.004

Lynn, C. W., & Bassett, D. S. (2019). The physics of brain network structure, function and control. Nature Reviews Physics, 1(5), 318–332. 10.1038/s42254-019-0040-8

Marcus, D., Harwell, J., Olsen, T., Hodge, M., Glasser, M., Prior, F., Jenkinson, M., Laumann, T., Curtiss, S., & Van Essen, D. (2011). Informatics and Data Mining Tools and Strategies for the Human Connectome Project [Technology Report]. Frontiers in Neuroinformatics, 5. https://www.frontiersin.org/journals/neuroinformatics/articles/10.3389/fninf.2011.00004

Margulies, D. S., Ghosh, S. S., Goulas, A., Falkiewicz, M., Huntenburg, J. M., Langs, G., Bezgin, G., Eickhoff, S. B., Castellanos, F. X., Petrides, M., Jefferies, E., & Smallwood, J. (2016). Situating the default-mode network along a principal gradient of macroscale cortical organization. Proc Natl Acad Sci U S A, 113(44), 12574–12579. 10.1073/pnas.1608282113

Maslov, S., & Sneppen, K. (2002). Specificity and Stability in Topology of Protein Networks. Science, 296(5569), 910–913. 10.1126/science.1065103

Medaglia, J. D., Huang, W., Karuza, E. A., Kelkar, A., Thompson-Schill, S. L., Ribeiro, A., & Bassett, D. S. (2018). Functional Alignment with Anatomical Networks is Associated with Cognitive Flexibility. Nat Hum Behav, 2(2), 156–164. 10.1038/s41562-017-0260-9

Mehta, K., Salo, T., Madison, T. J., Adebimpe, A., Bassett, D. S., Bertolero, M., Cieslak, M., Covitz, S., Houghton, A., Keller, A. S., Lundquist, J. T., Luo, A., Miranda-Dominguez, O., Nelson, S. M., Shafiei, G., Shanmugan, S., Shinohara, R. T., Smyser, C. D., Sydnor, V. J., … Satterthwaite, T. D. (2024). XCP-D: A Robust Pipeline for the Post-Processing of fMRI data. Imaging Neuroscience. 10.1162/imag_a_00257

Misic, B., Betzel, R. F., de Reus, M. A., van den Heuvel, M. P., Berman, M. G., McIntosh, A. R., & Sporns, O. (2016). Network-Level Structure-Function Relationships in Human Neocortex. Cereb Cortex, 26(7), 3285–3296. 10.1093/cercor/bhw089

Misic, B., Betzel, R. F., Nematzadeh, A., Goni, J., Griffa, A., Hagmann, P., Flammini, A., Ahn, Y. Y., & Sporns, O. (2015). Cooperative and Competitive Spreading Dynamics on the Human Connectome. Neuron, 86(6), 1518–1529. 10.1016/j.neuron.2015.05.035

Mueller, S., Wang, D., Fox, M. D., Yeo, B. T., Sepulcre, J., Sabuncu, M. R., Shafee, R., Lu, J., & Liu, H. (2013). Individual variability in functional connectivity architecture of the human brain. Neuron, 77(3), 586–595. 10.1016/j.neuron.2012.12.028

Murray, J. D., Bernacchia, A., Freedman, D. J., Romo, R., Wallis, J. D., Cai, X., Padoa-Schioppa, C., Pasternak, T., Seo, H., Lee, D., & Wang, X. J. (2014). A hierarchy of intrinsic timescales across primate cortex. Nat Neurosci, 17(12), 1661–1663. 10.1038/nn.3862

Neudorf, J., Kress, S., & Borowsky, R. (2022). Structure can predict function in the human brain: a graph neural network deep learning model of functional connectivity and centrality based on structural connectivity. Brain Structure and Function, 227(1), 331–343. 10.1007/s00429-021-02403-8

Raut, R. V., Snyder, A. Z., & Raichle, M. E. (2020). Hierarchical dynamics as a macroscopic organizing principle of the human brain. Proc Natl Acad Sci U S A, 117(34), 20890–20897. 10.1073/pnas.2003383117

Roberts, J. A., Perry, A., Roberts, G., Mitchell, P. B., & Breakspear, M. (2017). Consistency-based thresholding of the human connectome. Neuroimage, 145(Pt A), 118–129. 10.1016/j.neuroimage.2016.09.053

Rubinov, M., & Sporns, O. (2010). Complex network measures of brain connectivity: Uses and interpretations. Neuroimage, 52(3), 1059–1069. 10.1016/j.neuroimage.2009.10.003

Sarwar, T., Tian, Y., Yeo, B. T. T., Ramamohanarao, K., & Zalesky, A. (2021). Structure-function coupling in the human connectome: A machine learning approach. Neuroimage, 226, 117609. 10.1016/j.neuroimage.2020.117609

Schaefer, A., Kong, R., Gordon, E. M., Laumann, T. O., Zuo, X.-N., Holmes, A. J., Eickhoff, S. B., & Yeo, B. T. T. (2018). Local-Global Parcellation of the Human Cerebral Cortex from Intrinsic Functional Connectivity MRI. Cerebral Cortex, 28(9), 3095–3114. 10.1093/cercor/bhx179

Seguin, C., Jedynak, M., David, O., Mansour, S., Sporns, O., & Zalesky, A. (2023). Communication dynamics in the human connectome shape the cortex-wide propagation of direct electrical stimulation. Neuron, 111(9), 1391–1401 e1395. 10.1016/j.neuron.2023.01.027

Seguin, C., Mansour, L. S., Sporns, O., Zalesky, A., & Calamante, F. (2022). Network communication models narrow the gap between the modular organization of structural and functional brain networks. Neuroimage, 257, 119323. 10.1016/j.neuroimage.2022.119323

Seguin, C., Sporns, O., & Zalesky, A. (2023). Brain network communication: concepts, models and applications. Nat Rev Neurosci. 10.1038/s41583-023-00718-5

Smith, R. E., Tournier, J.-D., Calamante, F., & Connelly, A. (2012). Anatomically-constrained tractography: improved diffusion MRI streamlines tractography through effective use of anatomical information. Neuroimage, 62(3), 1924–1938. 10.1016/j.neuroimage.2012.06.005

Smith, R. E., Tournier, J.-D., Calamante, F., & Connelly, A. (2015). SIFT2: Enabling dense quantitative assessment of brain white matter connectivity using streamlines tractography. Neuroimage, 119, 338–351. 10.1016/j.neuroimage.2015.06.092

Smith, S. M., Jenkinson, M., Woolrich, M. W., Beckmann, C. F., Behrens, T. E. J., Johansen-Berg, H., Bannister, P. R., De Luca, M., Drobnjak, I., & Flitney, D. E. (2004). Advances in functional and structural MR image analysis and implementation as FSL. Neuroimage, 23, S208–S219.

Smolders, L., De Baene, W., Rutten, G.-J., van der Hofstad, R., & Florack, L. (2024). Can structure predict function at individual level in the human connectome? Brain Structure and Function, 229(5), 1209–1223. 10.1007/s00429-024-02796-2

Soman, S. M., Vijayakumar, N., Thomson, P., Ball, G., Hyde, C., & Silk, T. J. (2023). Cortical structural and functional coupling during development and implications for attention deficit hyperactivity disorder. Transl Psychiatry, 13(1), 252. 10.1038/s41398-023-02546-8

Somerville, L. H., Bookheimer, S. Y., Buckner, R. L., Burgess, G. C., Curtiss, S. W., Dapretto, M., Elam, J. S., Gaffrey, M. S., Harms, M. P., Hodge, C., Kandala, S., Kastman, E. K., Nichols, T. E., Schlaggar, B. L., Smith, S. M., Thomas, K. M., Yacoub, E., Van Essen, D. C., & Barch, D. M. (2018). The Lifespan Human Connectome Project in Development: A large-scale study of brain connectivity development in 5-21 year olds. Neuroimage, 183, 456–468. 10.1016/j.neuroimage.2018.08.050

Sporns, O. (2011). The human connectome: a complex network. Ann N Y Acad Sci, 1224(1), 109–125. 10.1111/j.1749-6632.2010.05888.x

Sporns, O., & Betzel, R. F. (2016). Modular Brain Networks. Annu Rev Psychol, 67, 613–640. 10.1146/annurev-psych-122414-033634

Suárez, L. E., Markello, R. D., Betzel, R. F., & Misic, B. (2020). Linking Structure and Function in Macroscale Brain Networks. Trends Cogn Sci, 24(4), 302–315. 10.1016/j.tics.2020.01.008

Suárez, L. E., Richards, B. A., Lajoie, G., & Misic, B. (2021). Learning function from structure in neuromorphic networks. Nature Machine Intelligence, 3(9), 771–786.

Sydnor, V. J., Cieslak, M., Duprat, R., Deluisi, J., Flounders, M. W., Long, H., Scully, M., Balderston, N. L., Sheline, Y. I., Bassett, D. S., Satterthwaite, T. D., & Oathes, D. J. (2022). Cortical-subcortical structural connections support transcranial magnetic stimulation engagement of the amygdala. Sci Adv, 8(25), eabn5803. 10.1126/sciadv.abn5803

Sydnor, V. J., Larsen, B., Bassett, D. S., Alexander-Bloch, A., Fair, D. A., Liston, C., Mackey, A. P., Milham, M. P., Pines, A., Roalf, D. R., Seidlitz, J., Xu, T., Raznahan, A., & Satterthwaite, T. D. (2021). Neurodevelopment of the association cortices: Patterns, mechanisms, and implications for psychopathology. Neuron, 109(18), 2820–2846. 10.1016/j.neuron.2021.06.016

Tournier, J.-D., Smith, R., Raffelt, D., Tabbara, R., Dhollander, T., Pietsch, M., Christiaens, D., Jeurissen, B., Yeh, C.-H., & Connelly, A. (2019). MRtrix3: A fast, flexible and open software framework for medical image processing and visualisation. Neuroimage, 202, 116137. 10.1016/j.neuroimage.2019.116137

Tustison, N. J., Avants, B. B., Cook, P. A., Zheng, Y., Egan, A., Yushkevich, P. A., & Gee, J. C. (2010). N4ITK: Improved N3 Bias Correction. IEEE Transactions on Medical Imaging, 29(6), 1310–1320. 10.1109/TMI.2010.2046908

Valk, S. L., Xu, T., Paquola, C., Park, B. Y., Bethlehem, R. A. I., Vos de Wael, R., Royer, J., Masouleh, S. K., Bayrak, S., Kochunov, P., Yeo, B. T. T., Margulies, D., Smallwood, J., Eickhoff, S. B., & Bernhardt, B. C. (2022). Genetic and phylogenetic uncoupling of structure and function in human transmodal cortex. Nat Commun, 13(1), 2341. 10.1038/s41467-022-29886-1

van den Heuvel, M. P., & Sporns, O. (2013). Network hubs in the human brain. Trends Cogn Sci, 17(12), 683–696. 10.1016/j.tics.2013.09.012

Van Essen, D. C., Smith, S. M., Barch, D. M., Behrens, T. E. J., Yacoub, E., & Ugurbil, K. (2013). The WU-Minn Human Connectome Project: An overview. Neuroimage, 80, 62–79. 10.1016/j.neuroimage.2013.05.041

Vasa, F., & Misic, B. (2022). Null models in network neuroscience. Nat Rev Neurosci. 10.1038/s41583-022-00601-9

Vazquez-Rodriguez, B., Suarez, L. E., Markello, R. D., Shafiei, G., Paquola, C., Hagmann, P., van den Heuvel, M. P., Bernhardt, B. C., Spreng, R. N., & Misic, B. (2019). Gradients of structure-function tethering across neocortex. Proc Natl Acad Sci U S A, 116(42), 21219–21227. 10.1073/pnas.1903403116

Veraart, J., Novikov, D. S., Christiaens, D., Ades-aron, B., Sijbers, J., & Fieremans, E. (2016). Denoising of diffusion MRI using random matrix theory. Neuroimage, 142, 394–406. 10.1016/j.neuroimage.2016.08.016

Wu, Z., Trevino, A. E., Wu, E., Swanson, K., Kim, H. J., D’Angio, H. B., Preska, R., Charville, G. W., Dalerba, P. D., Egloff, A. M., Uppaluri, R., Duvvuri, U., Mayer, A. T., & Zou, J. (2022). Graph deep learning for the characterization of tumour microenvironments from spatial protein profiles in tissue specimens. Nat Biomed Eng, 6(12), 1435–1448. 10.1038/s41551-022-00951-w

Yeo, B. T. T., Krienen, F. M., Sepulcre, J., Sabuncu, M. R., Lashkari, D., Hollinshead, M., Roffman, J. L., Smoller, J. W., Zöllei, L., & Polimeni, J. R. (2011). The organization of the human cerebral cortex estimated by intrinsic functional connectivity. Journal of Neurophysiology.

You, J., Ying, R., & Leskovec, J. (2021). Design Space for Graph Neural Networks. 10.48550/arXiv.2011.08843

Zalesky, A., Sarwar, T., Tian, Y., Liu, Y., Yeo, B. T. T., & Ramamohanarao, K. (2024). Predicting an individual’s functional connectivity from their structural connectome: Evaluation of evidence, recommendations and future prospects. Network Neuroscience, 1–32. 10.1162/netn_a_00400

Zamani Esfahlani, F., Faskowitz, J., Slack, J., Misic, B., & Betzel, R. F. (2022). Local structure-function relationships in human brain networks across the lifespan. Nat Commun, 13(1), 2053. 10.1038/s41467-022-29770-y

Zarkali, A., McColgan, P., Leyland, L. A., Lees, A. J., Rees, G., & Weil, R. S. (2021). Organisational and neuromodulatory underpinnings of structural-functional connectivity decoupling in patients with Parkinson’s disease. Commun Biol, 4(1), 86. 10.1038/s42003-020-01622-9

Zhou, J., Cui, G., Hu, S., Zhang, Z., Yang, C., Liu, Z., Wang, L., Li, C., & Sun, M. (2020). Graph neural networks: A review of methods and applications. AI open, 1, 57–81. 10.1016/j.aiopen.2021.01.001

Zimmermann, J., Griffiths, J., Schirner, M., Ritter, P., & McIntosh, A. R. (2019). Subject specificity of the correlation between large-scale structural and functional connectivity. Netw Neurosci, 3(1), 90–106. 10.1162/netn_a_00055

