## Supplementary Figures for "Group-common and individual-specific effects of structure-function coupling in human brain networks with graph neural networks"

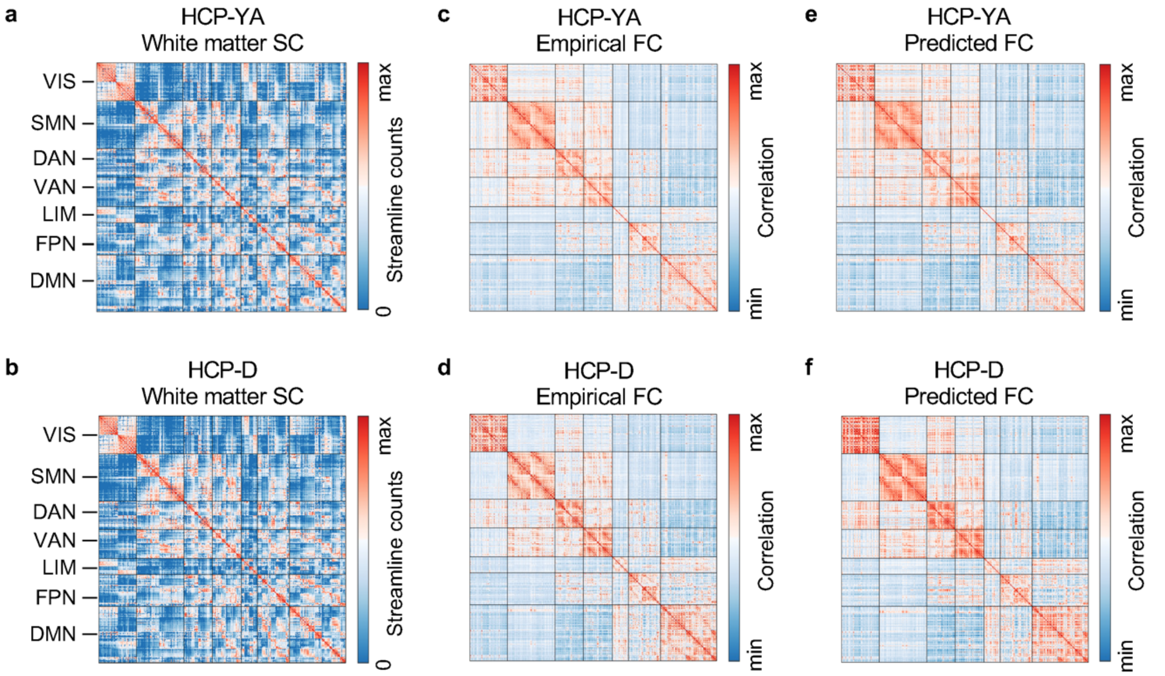


**Fig. S1. The group-averaged matrices of structural connectivity, empirical functional connectivity, and predicted functional connectivity from GNN. a**,**b**, The white matter structural connectivity (SC) in the HCP-YA (**a**) and HCP-D (**b**) datasets. **c**,**d**, The empirical functional connectivity (FC) in the HCP-YA (**c**) and HCP-D (**d**) datasets. **e**,**f**, The predicted FC in the HCP-YA (**e**) and HCP-D (**f**) datasets. GNN, graph neural network; HCP-YA, HCP young adult; HCP-D, HCP development; VIS, visual network; SMN, somatomotor network; DAN, dorsal attention network; VAN, ventral attention network; LIM, limbic network; FPN, frontoparietal network; DMN, default mode network.

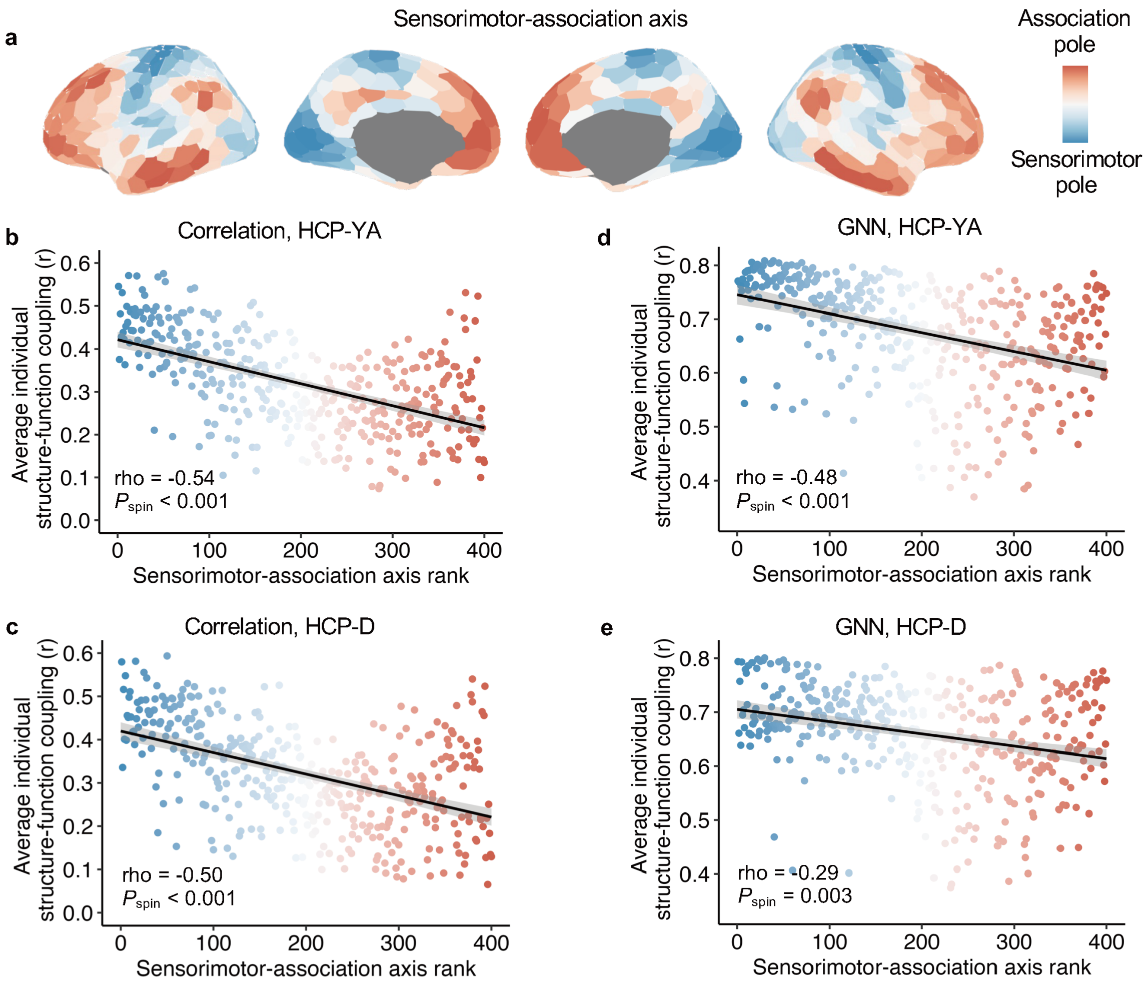


**Fig. S2. Regional structure-function coupling is distributed along sensorimotor-association axis across the cortex. a**, The cortical map of the sensorimotor-association axis was derived from Sydnor et al. (2021), where warm colors represent association cortices with higher ranks and cold colors represent sensorimotor cortices with lower ranks. **b**,**c**, The group-average structure-function coupling using Pearson correlation (**Fig. 2g** & **2h**) was negatively correlated with the sensorimotor-association axis ranks across all cortical regions in both HCP-YA (**b**, Spearman's rho = -0.54, $P_{spin}$ < 0.001) and HCP-D (**c**, Spearman's rho = -0.50, $P_{spin}$ < 0.001) datasets. **d**,**e**, The group-average GNN-based structure-function coupling (**Fig. 2i** & **2j**) was negatively correlated with the sensorimotor-association axis ranks across all cortical regions in the HCP-YA (**d**, Spearman's rho = -0.48, $P_{spin}$ < 0.001) and HCP-D (**e**, Spearman's rho = -0.29, $P_{spin}$ = 0.003) datasets. The outliers (mean ± 3×SD) were excluded in the correlation analysis. Each point on the scatter plot represents a cortical region and is colored by its sensorimotor-association axis rank. GNN, graph neural network; SD, standard deviation.

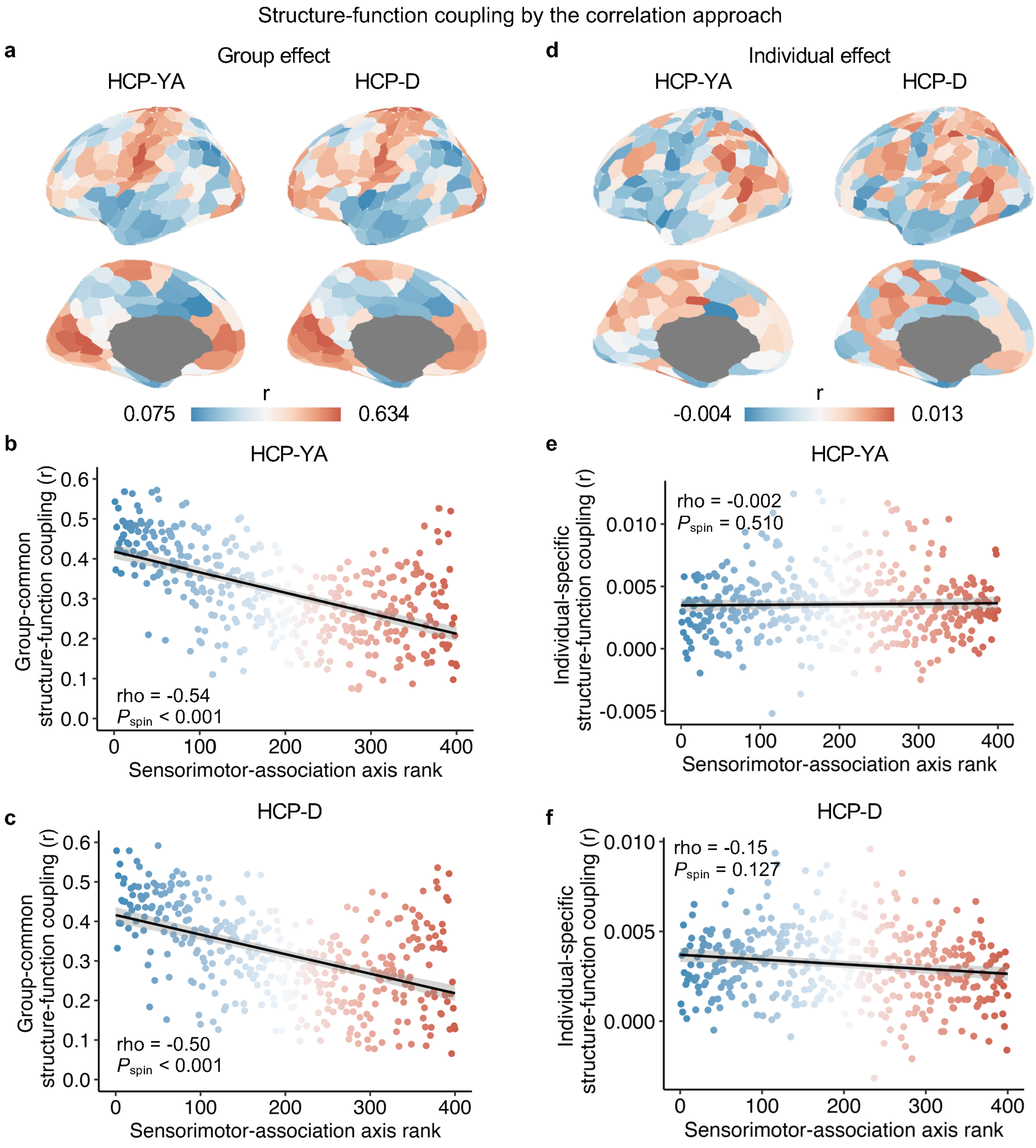


**Fig. S3. Group-common and individual-specific structure-function effects estimated using the correlation approach. a**, The cortical map of group-common coupling effects estimated by the Pearson correlation in the HCP-YA and HCP-D datasets. **b**,**c**, The group-common coupling effects were negatively correlated with the sensorimotor-association axis rank in the HCP-YA (**c**, Spearman's rho = -0.54, $P_{spin}$ < 0.001) and HCP-D (**d**, Spearman's rho = -0.50, $P_{spin}$ < 0.001) datasets. **d**, The cortical map of individual-specific coupling effects estimated by the Pearson correlation in the HCP-YA and HCP-D datasets. **e**,**f**, The individual-specific coupling effects were not significantly correlated with the sensorimotor-association axis rank in the HCP-YA (**e**, Spearman's rho = -0.002, $P_{spin}$ = 0.510) and HCP-D (**f**, Spearman's rho = -0.15, $P_{spin}$ = 0.127) datasets. The outliers (mean ± 3×SD) were excluded in the correlation analysis. Each point on the scatter plot represents a cortical region and is colored by its sensorimotor-association axis rank. SD, standard deviation.

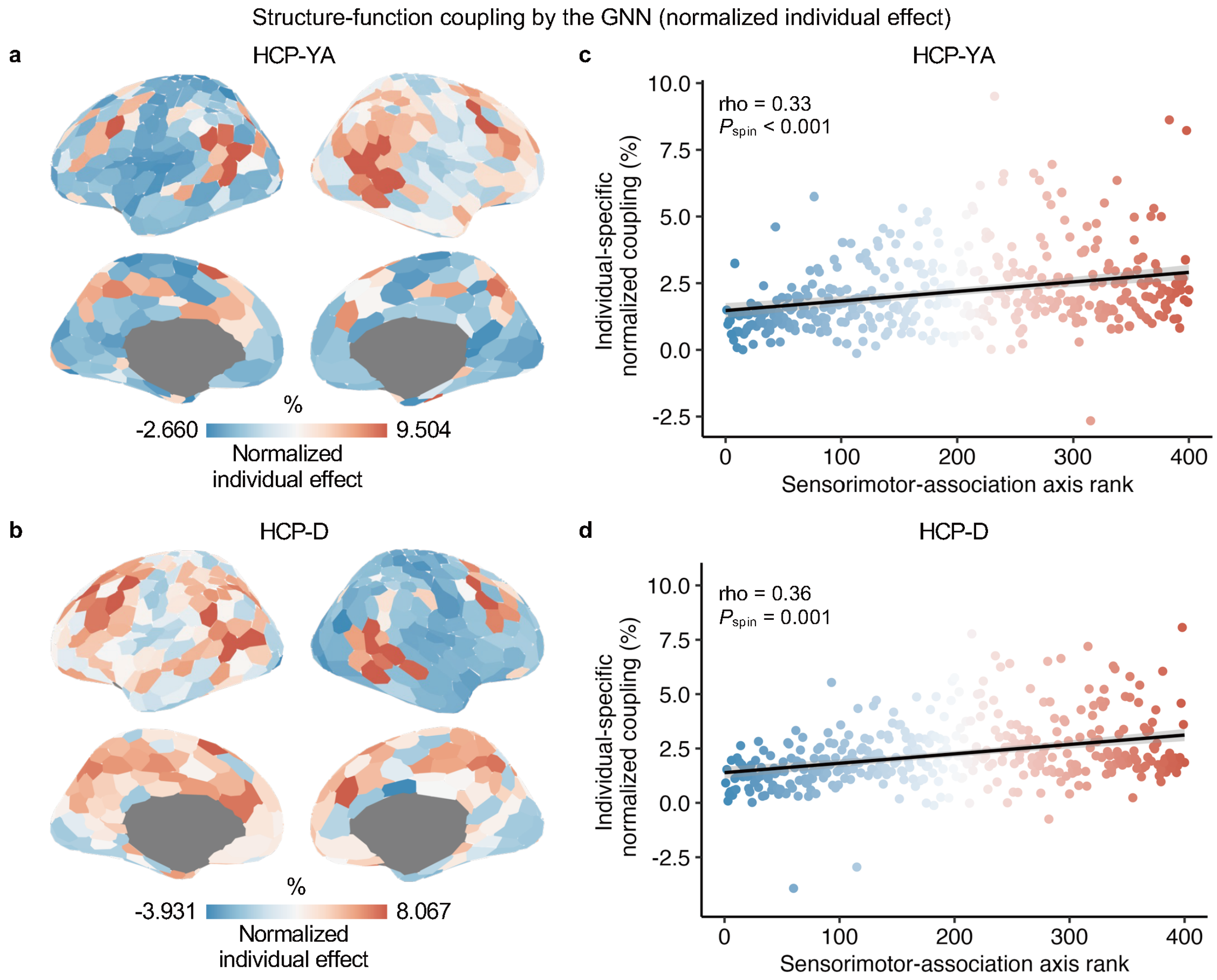


**Fig. S4. Normalized individual-specific structure-function effects align with the sensorimotor-association axis. a**,**b**, The cortical map of normalized individual-specific coupling effects (% of the total effects) estimated by the GNN model in the HCP-YA (**a**) and HCP-D (**b**) datasets. **c**,**d**, The normalized individual-specific coupling effects were positively correlated with the sensorimotor-association axis rank in the HCP-YA (**c**, Spearman's rho = 0.33, $P_{spin} < 0.001$) and HCP-D (**d**, Spearman's rho = 0.36, $P_{spin} = 0.001$) datasets. The outliers (mean ± 3×SD) were excluded in the correlation analysis. Each point on the scatter plot represents a cortical region and is colored by its sensorimotor-association axis rank. GNN, graph neural network; SD, standard deviation.

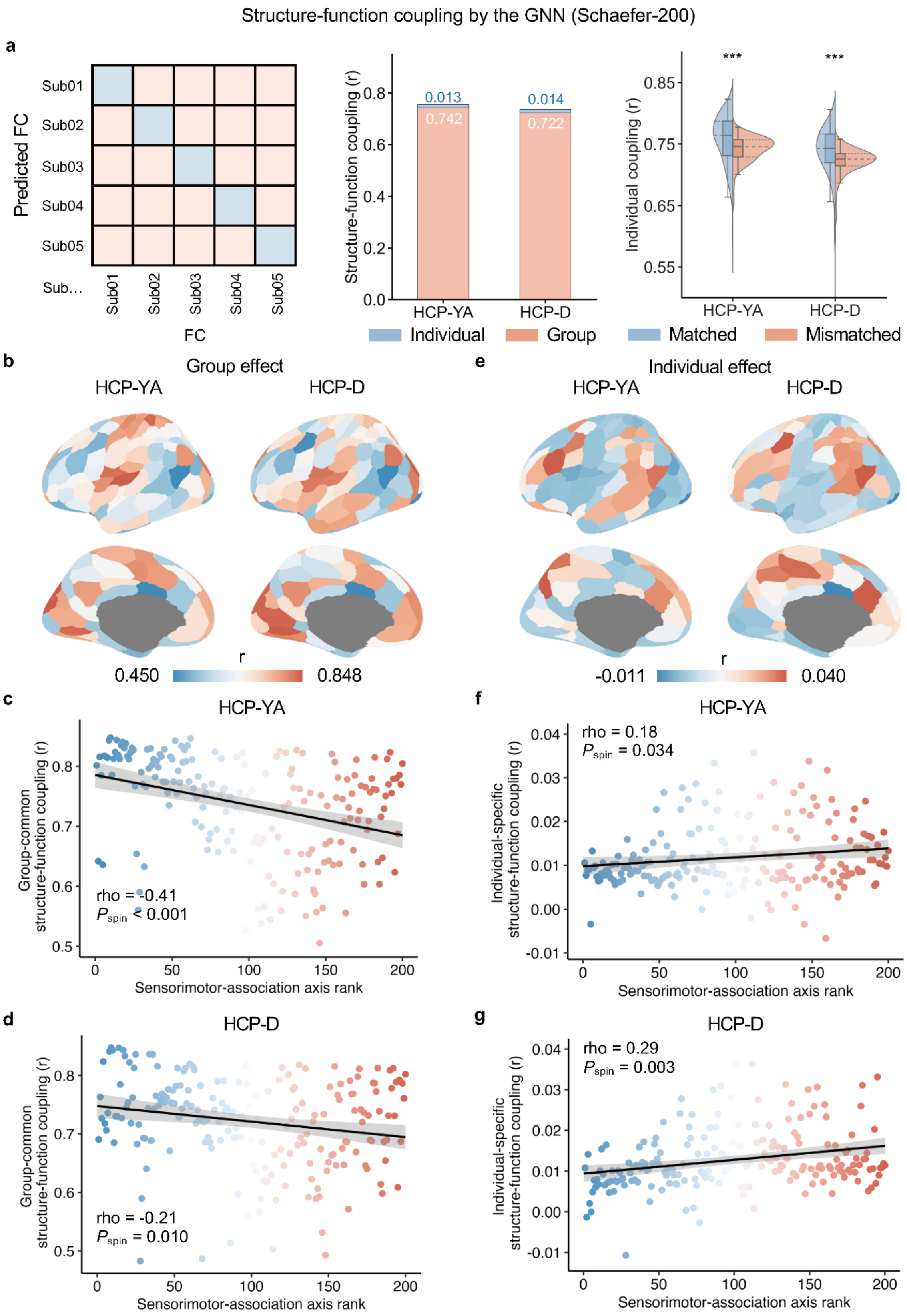

Structure-function coupling by the GNN (Schaefer-200)
a
Predicted FC
Sub01
Sub02
Sub03
Sub04
Sub05
Sub…
FC
Structure-function coupling (r)
0.013
0.742
0.014
0.722
HCP-YA
HCP-D
Individual
Group
Individual coupling (r)
***
***
Matched
Mismatched
b
Group effect
HCP-YA
HCP-D
r
0.450
0.848
e
Individual effect
HCP-YA
HCP-D
r
-0.011
0.040
c
HCP-YA
Group-common structure-function coupling (r)
rho = -0.41
Pspin < 0.001
Sensorimotor-association axis rank
d
HCP-D
rho = -0.21
Pspin = 0.010
f
HCP-YA
Individual-specific structure-function coupling (r)
rho = 0.18
Pspin = 0.034
g
HCP-D
rho = 0.29
Pspin = 0.003

**Fig. S5. GNN-derived group-common and individual-specific effects using Schaefer-200 cortical parcellation.** **a**, The structure-function coupling matrix between each pair of participants. Diagonal elements (cold color) represent the within-subject coupling (i.e., matched coupling), while off-diagonal elements (warm color) represent the between-subject coupling (i.e., mismatched coupling). The average value of the off-diagonal elements is defined as the group effect of structure-function coupling. The individual effect structure-function coupling is defined as the value of diagonal over non-diagonal. The group and individual effects were estimated using the Schaefer-200 (HCP-YA: 0.742/0.013; HCP-D: 0.722/0.014). For each participant, the matched coupling and average mismatched coupling were calculated and compared. The matched individual coupling values were significantly higher than the mismatched coupling. **b**, The cortical map of group-common effects using Schaefer-200 parcellations for the HCP-YA and HCP-D datasets. **c**,**d**, With Schaefer-200 atlas, the group-common effects were negatively correlated with the sensorimotor-association axis rank in the HCP-YA (**c**, Spearman's rho = -0.41, $P_{\text{spin}} < 0.001$) and HCP-D (**d**, Spearman's rho = -0.21, $P_{\text{spin}} = 0.010$) datasets. **e**, The cortical map of individual-specific effects using Schaefer-200 parcellations for the HCP-YA and HCP-D datasets. **f**,**g**, With Schaefer-200 atlas, the individual-specific coupling effects were positively correlated with the sensorimotor-association axis rank in the HCP-YA (**f**, Spearman's rho = 0.18, $P_{\text{spin}} = 0.034$) and HCP-D (**g**, Spearman's rho = 0.29, $P_{\text{spin}} = 0.003$) datasets. *** indicates $P < 0.0001$, two-tailed paired t-test. The outliers (mean ± 3×SD) were excluded in the correlation analysis. Each point on the scatter plot represents a cortical region and is colored by its sensorimotor-association axis rank. GNN, graph neural network; SD, standard deviation.

### References

Sydnor, V. J., Larsen, B., Bassett, D. S., Alexander-Bloch, A., Fair, D. A., Liston, C., Mackey, A. P., Milham, M. P., Pines, A., Roalf, D. R., Seidlitz, J., Xu, T., Raznahan, A., & Satterthwaite, T. D. (2021). Neurodevelopment of the association cortices: Patterns, mechanisms, and implications for psychopathology. *Neuron*, *109*(18), 2820-2846. https://doi.org/10.1016/j.neuron.2021.06.016
